# Regeneration recapitulates many embryonic processes, including reuse of developmental regulatory regions

**DOI:** 10.1101/2024.07.04.601589

**Authors:** Kaitlyn Loubet-Senear, Mansi Srivastava

## Abstract

The wide distribution of regenerative capacity across the animal tree of life raises the question of how regeneration has evolved in distantly-related animals. Given that whole-body regeneration shares the same end-point – formation of a functional body plan – as embryonic development, it has been proposed that regeneration likely recapitulates developmental processes to some extent. Therefore, understanding how developmental processes are reactivated during regeneration is important for uncovering the evolutionary history of regeneration. Comparative transcriptomic studies in some species have revealed shared gene expression between development and regeneration, but it is not known whether these shared expression profiles correspond to shared functions, and which mechanisms activate expression of developmental genes during regeneration. We sought to address these questions using the acoel *Hofstenia miamia*, which is amenable to studies of both embryonic development and whole-body regeneration. By examining functionally validated regeneration processes during development at single-cell resolution, we found that whereas patterning and cellular differentiation are largely similar, wound response programs have distinct dynamics between development and regeneration. Chromatin accessibility analyses revealed that regardless of playing concordant or divergent roles during regeneration and development, genes expressed in both processes are frequently controlled by the same regulatory regions, potentially via utilization of distinct transcription factor binding sites. This study extends the known correspondence of development and regeneration from broad transcriptomic similarity to include patterning and differentiation processes. Further, our work provides a catalog of regulatory regions and binding sites that potentially regulate developmental genes during regeneration, fueling comparative studies of regeneration.

## INTRODUCTION

During animal development, tightly orchestrated processes act to establish an animal body, with diverse cell types, tissues, and organs formed precisely relative to each other to enable organismal function. In animal lineages capable of whole-body regeneration, the same end-point can be achieved post-embryonically – entire body plans and all missing cell types are restored upon injury in adults^1,2^. Thus, the extent to which regenerative pathways recapitulate developmental processes has been a long-standing question in the field of regenerative biology^3^. Given regenerative processes must recreate structures that had already been made once during development, one might expect a strong correspondence of molecular and cellular mechanisms between development and regeneration. Alternatively, given that the starting points of the two processes are quite different, with development beginning from a fertilized zygote and regeneration from a multicellular, complex remnant of an animal, one might expect some mechanisms to be distinct. Studies comparing development and regeneration in regenerative species could contribute to the understanding of the relationship between development and regeneration, and identify the mechanisms that enable the utilization of developmental pathways in adult regeneration, shedding light on how regenerative capacity evolved across animals.

Regeneration and development have been compared using a multitude of approaches in many species, ranging from focused studies of limbs and organs in vertebrates to unbiased comparisons using transcriptome profiling in invertebrates^4–17^. While cross-transplantation of developing limb buds and regenerating limb blastemas of axolotls suggested broadly shared mechanisms for the two processes^13^, more recent studies suggest that cell plasticity, the dependence on nerve signaling and immune cells, and even spatial expression of patterning genes in regenerating limbs are distinct from those in developing limbs in salamanders and other species^8,12,18^. Further, whereas transcriptome profiling in bulk shows broad redeployment of developmental genes during regeneration in many invertebrates^9–11,17^, differences have been reported in terms of order of tissue reconstruction (in hemichordates^17^), source of progenitor cells (in larval sea stars^16^), temporal order of gene expression (in crustaceans^9^), and signaling pathway functions (in sea anemones^11^). These studies leave open the question of how broad transcriptome-wide similarities between the processes of development and regeneration can be reconciled with the various reported differences, which are all in different organismal and mechanistic contexts.

Transcriptomic similarities obtained from tissues in bulk could be indicative of gene expression programs that operate in the same cellular contexts and contribute to the same functions in development and regeneration. Alternatively, these similarities could be obfuscating expression of genes in different cell types and their functions in new contexts, a scenario that would indicate regeneration co-opts developmental genes into new networks. We sought to explicitly examine transcriptome-wide gene expression during development and regeneration in a system that would enable an assessment of cellular context and regulatory relationships. We therefore focused on *Hofstenia miamia (H.miamia)*, an acoel worm that has recently been established as a new model organism for studying both development and regeneration^19,20^ (Figure 1A, 1B). Notably, *H. miamia* is capable of whole-body regeneration and also produces accessible embryos^21^. In addition to the availability of transcriptomic data from tissues in bulk over the course of development and regeneration, these processes have been sampled at single-cell resolution in *H. miamia*^22–25^(Figure 1B). Furthermore, this system is amenable to construction of gene regulatory networks, with chromatin profiling data available for both regeneration and development^22,26^. Therefore, this system is ideally poised for rigorous comparisons of cellular contexts and regulatory linkages during development and regeneration at transcriptome- and genome-wide scales.

**Figure 1.**
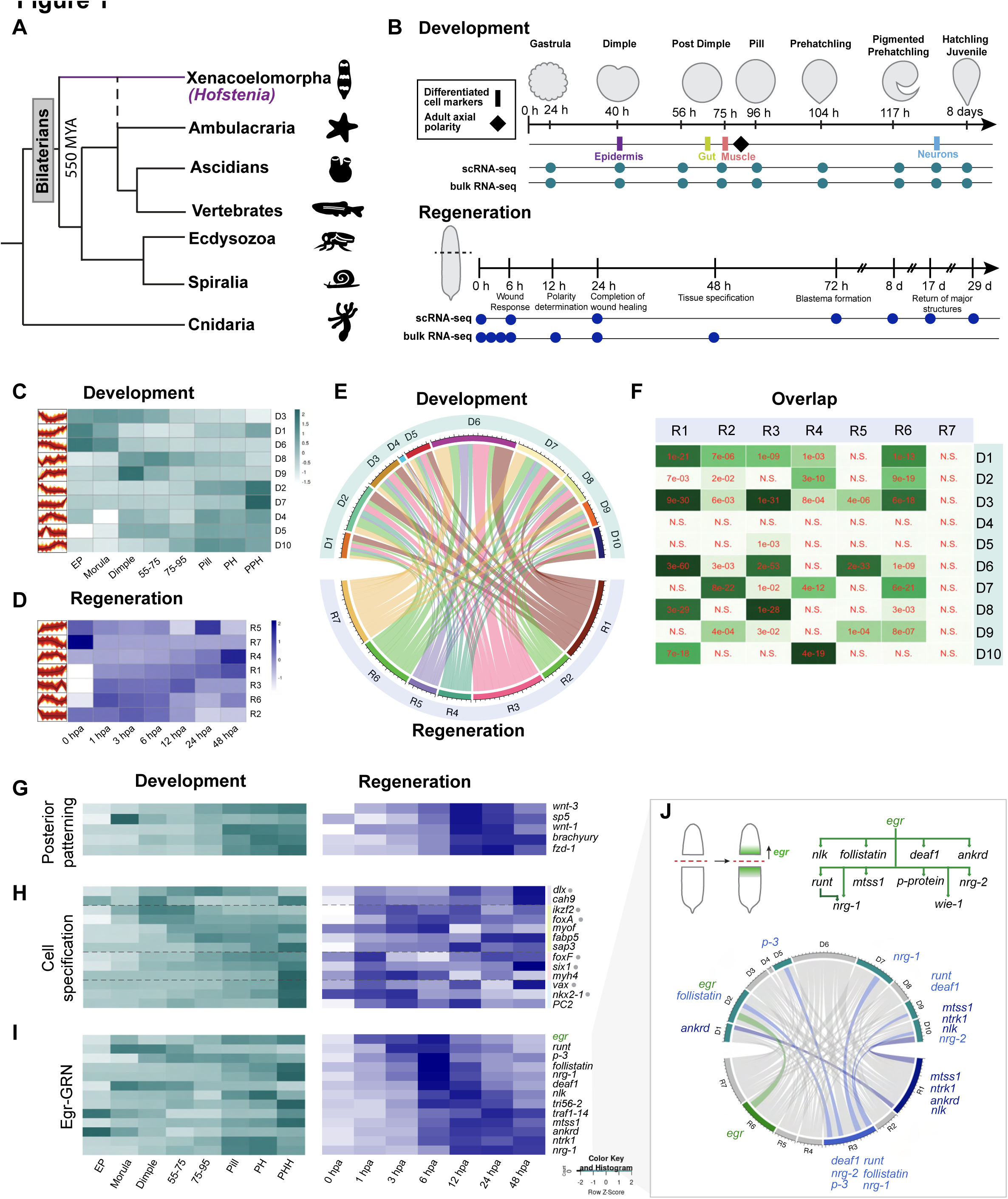
Developmental genes are redeployed during *H. miamia* regeneration in a variety of temporal patterns, some suggestive of shared modules and others consistent with distinct networks. A) Phylogenetic placement of *Hofstenia miamia* as either the outgroup of bilaterians or the sister to Ambulacraria (dashed line). B) Existing developmental and regenerative single-cell and bulk RNA-seq datasets. Developmental bulk: Kimura 2021^24^; Developmental single-cell: Kimura et al. 2022^25^; Regeneration bulk: Gehrke et al. 2019^22^; Regeneration single cell: Hulett et al. 2023^23^. Timing of detectable differentiated markers for major *H. miamia* tissue types (rectangles) and adult axial polarity (diamond) during development are indicated. Major regenerative hallmarks are noted. Time in hours (h) or days (d); development: this refers to time post post lay; regeneration: this refers to time post amputation. C) Fuzzy c-means clustering of developmental bulk RNA-seq data. Clusters are labeled D1-D10. Cluster profiles are shown next to the heatmap for cluster centroid values (shown in black line). D) Fuzzy c-means clustering of regenerative bulk RNA-seq data. Clusters are labeled R1-R7. Cluster profiles are shown next to the heatmap for cluster centroid values (shown in black line). E) Chord plot of overlapping genes in fuzzy clusters between development (top, D1-D10) and regeneration (bottom, R1-R7). Chord thickness is scaled based on number of genes overlapping between each pair of clusters. F) Gene overlap plot showing clusters with numbers of genes overlapping beyond expected by random chance. G-I) Bulk RNA-seq heatmaps for development and regeneration of candidate networks from *H. miamia* regeneration. EP – early pool; PH – prehatchling; PPH – pigmented prehatchling. G) Posterior patterning network, H) cell specification. Gray circles indicate transcription factors, genes associated with a given lineage based on Hulett et al. 2023^23^ are delineated by a dashed gray line. Purple represents epidermal genes, green are endodermal genes, salmon are muscle genes, and light blue are neural genes. I) Egr-GRN. J) Top: Egr-GRN. Following amputation *egr* is expressed at both wound sites and is required for expression of downstream genes. Bottom: Chord plot demonstrating correspondence of developmental and regenerative fuzzy clusters that contain genes from the Egr-GRN. Developmental clusters containing these genes are highlighted teal; regenerative clusters are highlighted in blue.

We began by comparing transcriptome profiles in bulk, and recapitulated findings of similar prior studies: many sets of *H. miamia* genes have similar expression dynamics as each other in both development and regeneration. Next, we leveraged our knowledge of three processes that have been functionally investigated during regeneration in *H. miamia* – wound response^22^, posterior patterning^27^, and cell fate specification^23,28^ – and found that although genes implicated in re-establishing the anterior-posterior axis in regeneration exhibit similar expression dynamics as each other in both development and regeneration, the temporal dynamics of genes involved in other characterized regenerative programs did not align between development and regeneration. By delving into these processes at single-cell resolution, we found that despite variable temporal patterns in bulk RNA-sequencing data, cells are likely specified using similar genetic factors during development and regeneration, and that differentiation proceeds along similar molecular trajectories. In contrast, the network architecture of a previously identified wound-induced program that deploys rapidly following injury appears to be a regeneration-specific innovation, as the temporal dynamics during development are inconsistent with the same regenerative network acting during development. Finally, we leveraged chromatin profiling data to identify the regulatory mechanisms that re-activate developmental genes during regeneration, and found that in many instances, including for the majority of wound-response genes, the same regulatory regions likely are needed for expression during both processes. We found that different binding sites could be utilized in these shared regulatory regions, potentially explaining distinct input mechanisms that lead to transcriptional activation of the same genes during development and regeneration. Our data suggest that the capacity to robustly regenerate likely relied upon the evolution of new regeneration-specific enhancers as well as the repurposing of developmental enhancers, consistent with recent data from other regenerative animals^4,15^. Comparing specific genetic networks and the evolution of these regulatory regions across animals will reveal the history of regeneration across the tree of animal life.

## RESULTS

### Developmental genes are redeployed during regeneration in patterns suggestive of both shared modules and distinct networks

We sought to assess the extent to which developmental genes are redeployed during regeneration. Using previously published bulk RNA-sequencing datasets from developing and regenerating *H. miamia*^22,24^, we identified sets of putatively coregulated gene by grouping transcripts with similar temporal expression dynamics, and then compared these gene sets between developmental and regeneration datasets (Figures 1B-F, see Materials and Methods). The developmental dataset spanned seven stages from early cleavage through hatching, and the regeneration dataset captured the first two days of regeneration (ranging from 0 - 48 hours post amputation, hpa), encompassing the wound response, re-establishment of patterning information, and specification of several major cell populations (Figure 1B)^22,23,27^. We used fuzzy c-means clustering to identify cohorts of genes with similar expression dynamics as each other during development and (separately) in regeneration, identifying suites of genes with peak expression at early, intermediate, and late stages of both processes (Figure 1C-D). Many of the rapidly increased genes during regeneration included those with previously characterized wound-induced expression^22^ (Supplemental Table 1), validating this method’s ability to group genes based on temporal expression.

We next assessed whether any of these cohorts of genes were shared between development and regeneration, which could indicate redeployment of developmental gene expression modules during regenerative processes. We found that each regenerative cohort contains subsets of genes from every developmental cohort (Figure 1E, F, Supplemental Figure 1A), suggesting that these gene subsets, visualized as ‘ribbons’ in the chord plot in Figure 1E, could be genetic modules or networks that mediate shared biological processes, albeit with distinct temporal patterns between regeneration and development. We therefore performed Gene ontology (GO) analysis, which revealed biological processes likely associated with these gene subsets (Supplemental Table 2). For example, genes in regeneration cohort 1 (R1), which are all upregulated upon amputation, include a gene subset that is highly expressed early in development (developmental cohort 6, D6) and is associated with apoptosis, as well as a gene subset that is expressed late in development (developmental cohort 10, D10) and is associated with Wnt signaling (Supplemental Figure 1B).

Next, to complement these transcriptome-wide comparisons, we performed more specific comparisons by focusing on three processes that had been previously studied functionally during regeneration in *H. miamia* – namely, posterior patterning, cell fate specification, and a wound-induced gene regulatory network under the control of the master regulator Egr (Egr-GRN)^22,23,27^ (Figures 1G-I, Supplemental Figure 1C). First, we found that genes previously implicated in a posterior patterning program during regeneration^27^ also exhibited similar temporal patterns as each other during development (Figure 1G), which is consistent with these genes operating together during development. Second, we considered cell fate specification. Functional studies during regeneration that focused on differentiation of neoblasts, the collectively pluripotent stem cells of *H. miamia*, had uncovered transcription factors (TFs) and differentiated marker genes involved in restoration of muscle, neural, digestive, and epidermal cell types^23^. These studies had also revealed that differentiation of stem cells into new tissues at the wound site first becomes detectable 48-72 hpa, and therefore we were unsurprised to see the lack of temporally coordinated gene expression in our bulk regeneration data, which spanned 0 - 48 hpa (Figure 1H). Furthermore, the existence of pre-existing differentiated tissues in regenerating fragments could dilute signal from newly differentiating tissue. However, despite the lack of shared temporal gene expression in regeneration data, these genes showed highly concordant developmental temporal expression profiles (Figure 1H). Moreover, markers for a given cell type began to be expressed during development at times consistent with the previously described developmental timing for each lineage^24^ (Figure 1A, 1H). Additionally, the onset of TF expression preceded that of its corresponding differentiated marker, consistent with the TFs controlling expression of these downstream genes in development as they do during regeneration (Figure 1H).

Finally, we investigated the Egr-GRN, which involves rapid transcriptional upregulation of the zinc finger TF Egr at wound sites in response to injury, with subsequent expression of other wound-induced Egr-target genes^22^ (Figure 1I,J). As expected, these genes show clearly coordinated expression 0-48 hpa. In contrast, when we investigated the developmental temporal dynamics of genes comprising this network, we did not detect any concordant expression of these genes relative to each other (Figure 1I). Notably, *egr* was not expressed until later developmental timepoints, after many of the other genes in this network have already become expressed. The architecture of this network downstream of Egr was also not intact during development – GRN member genes are contained within two regeneration cohorts (R1 and R3) but during development they are present in multiple cohorts with distinct temporal dynamics (Figure 1J). This suggests that the Egr-GRN is not operational during development.

Altogether, comparing development and regeneration at the level of bulk RNA-seq revealed indications both for shared (posterior patterning and cell fate specification) and divergent (Egr-GRN) pathways between the two processes. However, while temporally coordinated gene expression is consistent with shared gene functions between development and regeneration, it does not rule out that genes may be active in different cell types and in different processes in development relative to in regeneration.

### Single-cell analysis suggests that cell specification programs are shared across development and regeneration in *H. miamia*

We reasoned that single-cell RNA-sequencing (scRNA-seq) data would allow us to interrogate gene expression within the context of cellular lineages, and thus better assess the significance of the temporal dynamics we observed for genes in the regenerative cell-specification network. During *H. miamia* regeneration, new tissues are produced via differentiation of a pool of collectively pluripotent stem cells (Figure 2A)^21,23^. First, we evaluated the transcriptome-wide similarity between developmental and regenerative lineages by utilizing existing *H. miamia* scRNA-seq datasets^23,25^. Previously, a scRNA-seq time course of *H. miamia* development had been analyzed using the trajectory inference tool URD to predict differentiation trajectories based on transcriptional similarities between cells^25,29^. An URD tree progresses from early to late time points in development; branches represent predicted cell lineages, and nodes/branching points represent predicted lineage divergence. In the *H. miamia* developmental URD tree, the latest time point is the hatched juvenile worm. Because *H. miamia* undergoes direct development, the cells and tissues in this stage correspond to tissues in the adult, allowing branches to be assigned to their corresponding adult tissue types based on known cell markers (Figure 2B)^21,23,24^.

**Figure 2.**
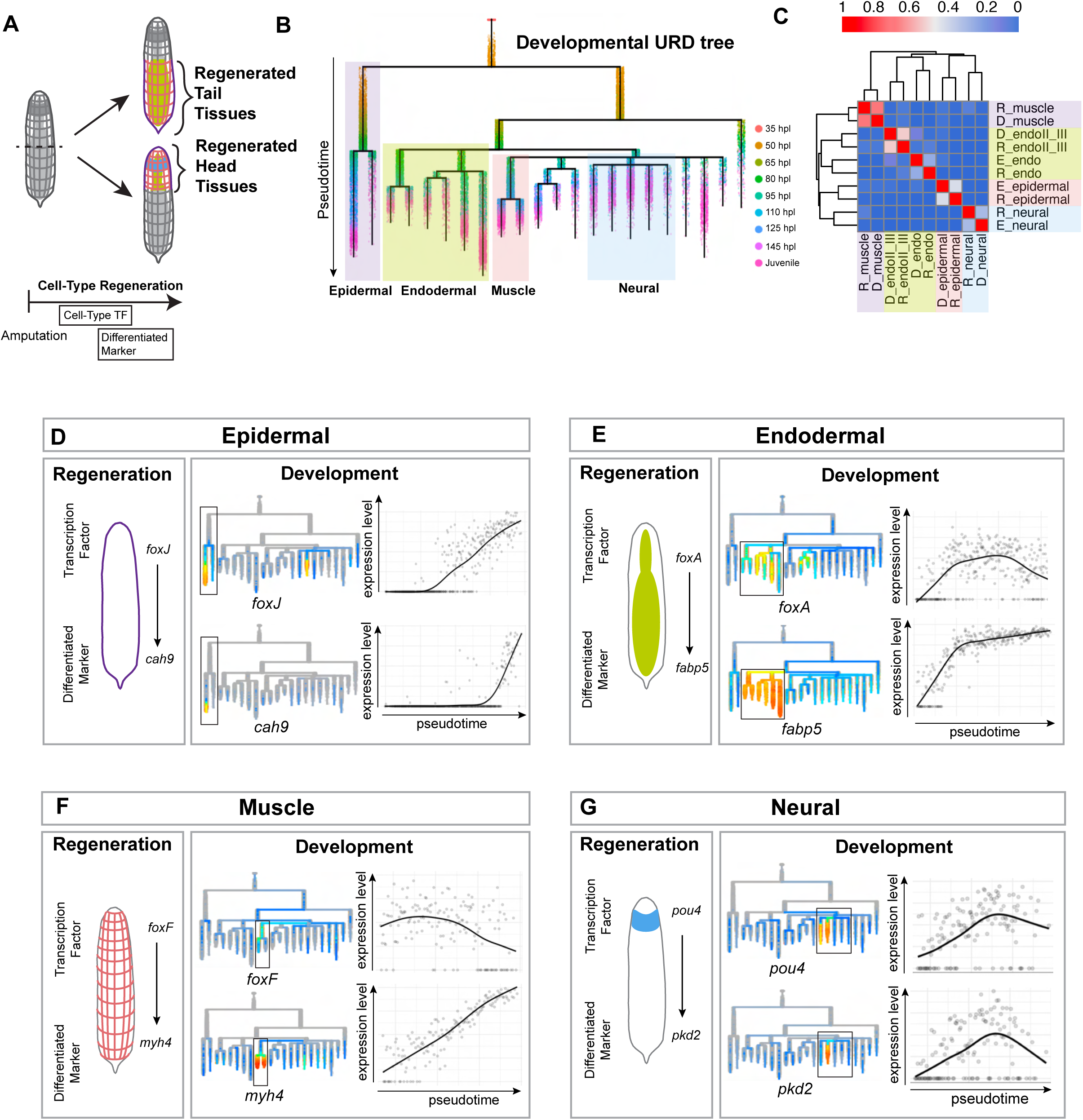
Single-cell analysis suggests that cell specification programs are shared across development and regeneration in *H. miamia*. A) *H. miamia* regenerates major cell populations following injury. Following amputation, cell-type transcription factors are expressed followed by re-emergence of differentiated markers in the regenerated tissue. B)*H. miamia* Developmental URD tree from Kimura, et al. 2022^25^ with major cell populations highlighted. Epidermal – purple, muscle – salmon, endodermal – green, neural – sky blue. Branches progress along pseudotime from the top to the bottom of the page. Cells from the earliest time point (35 hpl, salmon) populate the root and hatched juvenile (pink cells) populate the tips with other timepoints in between. C) Pearson correlation comparing pseudobulk gene expression of developmental and regenerative lineages. Red: more similar, blue: less similar. D-G) Transcription factors and differentiated markers from regeneration (left) for major cell populations (epidermal, endodermal, muscle, and neural) are associated with those same lineages in the developmental URD tree (boxed branches, middle). Spline plots show average expression of TFs and differentiated markers for these cell types within their developmental lineages (right).

To compare regenerative and developmental lineages, we extracted cells from lineages corresponding to each major tissue in the developmental URD tree (Figure 2B -lineages are boxed and colored), created pseudobulk gene expression matrices for these lineages to identify broad patterns of gene expression within each cell type, and calculated the Pearson correlation between these lineages and pseudobulk matrices of major regenerative cell types based on previous clustering^21,23,24^. We found that developmental cell populations (epidermal, endodermal, muscle, and neural) cluster with their corresponding regenerative cell population, suggesting at a transcriptome-wide level lineages are similar between development and regeneration (Figure 2C).

Given this broad lineage correspondence, we next used these URD-predicted lineages to query the developmental expression dynamics of TFs functionally implicated in regenerative production of cell populations. The bulk RNA-seq data had suggested that during development, expression of TFs specific for a given lineage precede expression of that lineage’s differentiated marker genes. Using the scRNA-seq data, we found that these genes are associated with the corresponding developmental lineage, and that within this differentiation trajectory TF expression preceded that of the differentiated marker, consistent with the patterns observed in the bulk dataset (Figure 2D, E, F, G, Supplemental Figure 2A). These data suggest that the regenerative differentiation trajectories are largely a re-use of developmental programs at a transcriptional level.

### Neural differentiation is similar in development and regeneration, but key regulators likely play additional non-neural roles during development

Given the broad similarities in lineage specification, we next investigated in more detail the specification of neural cell populations during development and regeneration, as this process had already been functionally interrogated during regeneration in *H. miamia*^28^. Notably, this regenerative network connects the early wound-response to specification of neural subpopulations, providing the opportunity to consider not only how cell populations are specified, but also compare how these trajectories might be initiated in development and regeneration. During regeneration of the anterior facing wound site (which will form the new head), wound-induced binding of the NFY transcription factor complex to its motifs leads to wound-induced expression of the neural progenitor marker *soxC*, which is required for expression of neural transcription factors that are subsequently required for the expression of differentiated neural subpopulation markers (Figure 3A).

**Figure 3.**
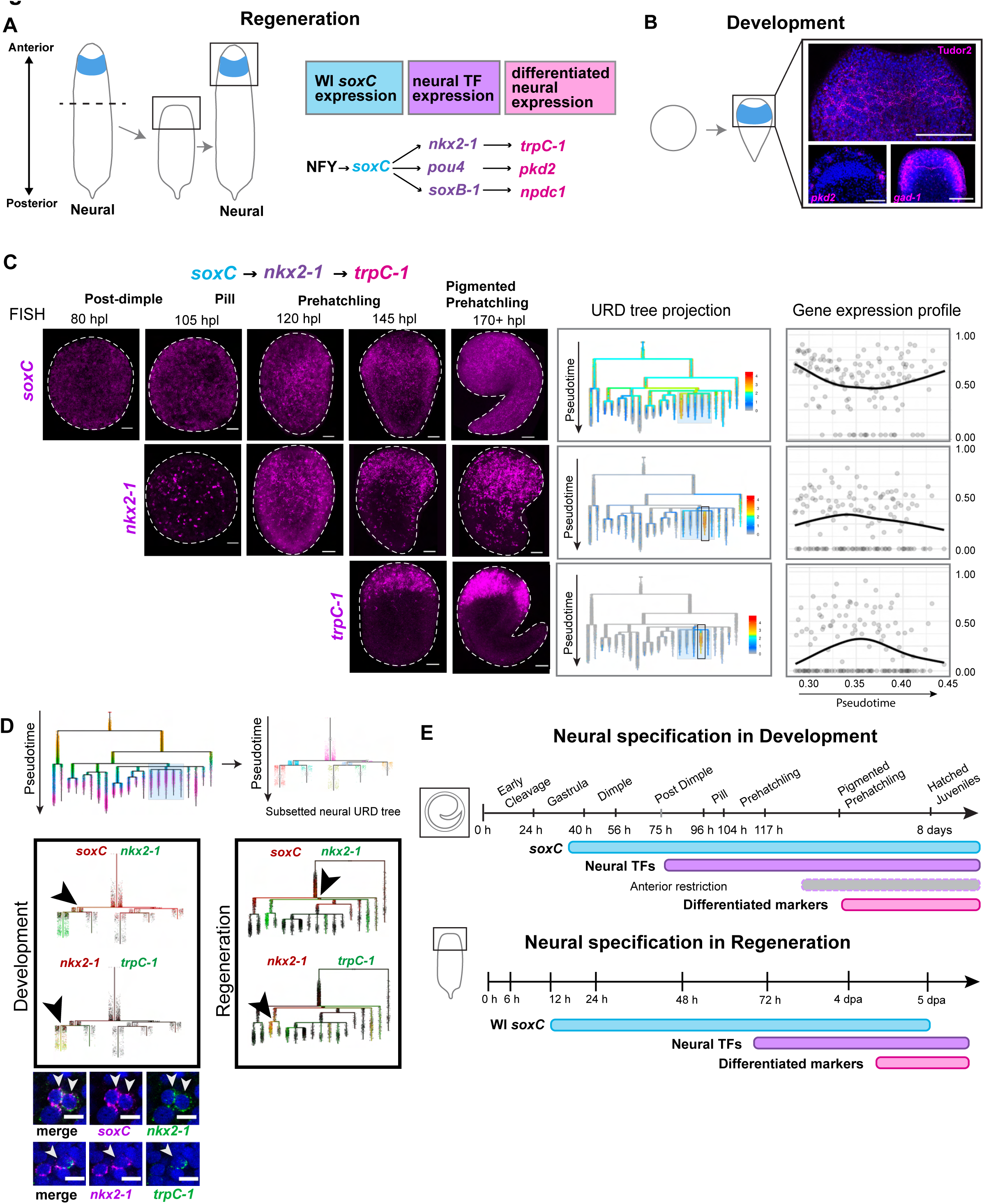
Neural differentiation proceeds via similar differentiation trajectories in development and regeneration, but key regulators likely play additional non-neural roles during development. A) Summary of neural cell population regeneration from Hulett, et al. 2023^28^. Following amputation, *H. miamia* tail fragments are capable of regenerating new brains at the anterior facing wound. NFY is required for wound-induced (WI) *soxC* expression (light blue), which is required for expression of neural transcription factors (light purple). These transcription factors are required for expression of differentiated neural markers (pink). B) Major neural structures, including the brain, are present upon hatching. Tudor is a custom *H. miamia* antibody that stains neurite bundles; *pkd2* marks a more external population of cells predicted to be sensory^28,63^, and *gad-1* shows the dorsal commissure^30^. 100um scale bars. C) Expression patterns of the *soxC > nkx2-1 > trpC-1* neural lineage during development. Left: Fluorescent in situ hybridization of *soxC, nkx2-1, and trpC-1* during development. Anterior is up. 50 um scale bars. Middle: Expression of neural genes on the developmental URD tree. Neural lineages are boxed. Right: Spline plot from URD tree showing average expression timing of neural genes across pseudotime corresponding to progression from URD tree root to tips. D) The developmental URD tree was subsetted to produce a more detailed neural URD tree. Regenerative Co-expression within branches of the URD tree was assessed for TF and downstream factor pairs in both development and regeneration: *soxC and nkx2-1,* and *nkx2-1 and trpC-1*. Co-expression of two transcripts within a cell on the URD tree is shown by yellow color; cells expressing only one gene or the other are red or dark green, and low expression of either transcript is black. E) Summary of regenerative neural gene expression during development (top, this study) and regeneration (bottom, Hulett, et al. 2023^28^). During development *soxC* is expressed from ∼35 hpl. Neural transcription factors exhibit expression starting at between 75 and 95 hpl (light purple), and then exhibit spatial restriction to hatched juvenile expression patterns around 120 hpl (gray), coincident with expression of differentiated markers (pink).

First, we confirmed that major cellular populations and features of the nervous system are present upon hatching. The *H. miamia* nervous system consists of an anterior condensation of neurons, which is detectable via multiple molecular markers, as well as a diffuse nerve net throughout the rest of the body^30^. We find all of these features are present upon hatching using a combination of immunohistochemistry (IHC) and fluorescent *in situ* hybridization (FISH) (Figure 3B, Supplemental Figure 3A). Next, we asked when these differentiated markers and their corresponding TFs, identified during regeneration, appear during development. Consistent with the previously reported expression for the differentiated neural marker *gad-1* during development^24^, markers of differentiated neural populations did not appear until late in development (∼145 hours post lay (hpl), during the pigmented prehatchling stage). The spatial expression pattern of these genes was similar at their onset in the pigmented prehatchling stage to that observed in hatched animals, and was consistent with labeling of neural populations (Figure 3C, Supplemental Figure 3B). This also corresponded to the emergence of the nerve net and anterior condensation of neurons based on IHC (Supplemental Figure 3C). When we looked at the developmental expression of neural TFs identified as playing a key role in the regeneration of neural cells (*nkx2-1, pou4, soxB-1)*, we found that their expression preceded corresponding differentiated markers that they target (Figure 3C, Supplemental Figure 3B).

However, unlike with the differentiated markers, they were initially expressed broadly and sparsely throughout the animal. Shortly preceding the onset of the differentiated neural markers the spatial expression of TFs took on the distinct patterns observed in the homeostatic animals: at later developmental time points *nkx2-1, soxB-1*, and *vax* were all enriched in the anterior, and *pou4* exhibited enrichment around the mouth and tip of the tail (Figure 3C, Supplemental Figure 3B). The neural progenitor marker *soxC* preceded expression of these neural progenitor markers, and exhibited broad expression throughout the embryo, similar to its homeostatic expression in hatched worms^28^ (Figure 3C). Overall, the timing of the expression onset of TFs and targets was consistent with the previously characterized regenerative differentiation trajectories.

Next we used scRNA-seq data to investigate whether these genes could be involved in similar developmental differentiation trajectories relative to the trajectories in regeneration. To determine more finely-resolved molecular trajectories, we subsetted cells associated with the neural lineages and generated an URD tree using these cells (Figure 3D). We found that in both the developmental and regenerative URD trees, neural TF/differentiated marker gene pairs were associated with the same lineages as each other. We next assessed whether these TF/differentiated marker pairs were co-expressed in predicted molecular trajectories: *soxC* was co-expressed with *nkx2-1, pou4,* and *soxB-1* (although the *soxC+/nkx2-1+, soxC+/pou4+* and *soxC+/soxB-1+*populations are distinct from each other). *nkx2-1* was then co-expressed with *trpC-1* in the URD tree, as *pou4* was with *pkd2*. (Figure 3D, Supplemental Figure 3D). Double FISH was consistent with these findings - there was co-expression between *soxC* and *nkx2-1,* and between *nkx2-1* and *trpC-1* (Figure 3D, Supplemental Figure 3E). Together, these data suggest that there is general concordance in specification of major cell populations in *H. miamia* development and regeneration, as well as for neuronal subpopulations.

However, the scRNA-seq data also showed that *soxC* was expressed outside of the neural lineage in earlier developmental timepoints, suggesting that *soxC* may have broader roles during development. The components of the NFY complex also exhibit broad cell type expression during development based on the scRNA-seq data, but unlike *soxC* we note that these genes are similarly broad in regeneration, making it difficult to assess based on expression alone what NFY’s developmental role might be (Supplemental Figure 3F). In other systems, NFY is known to act both as a pioneer transcription factor to increase chromatin accessibility but also to have roles in tissue-specific gene regulation^24,31^. We noted that the difference in cellular contexts between development and regeneration expanded to other wound-induced genes, where often genes in the Egr-GRN either showed expanded or restricted developmental expression relative to regeneration (Supplemental Figure 3G). Overall, the timing we observe with neural genes during development is consistent with developmental differentiation trajectories matching the regenerative differentiation, with potential additional developmental roles for the early network components *soxC* and NFY.

### Spatial and cell-type expression suggest shared posterior patterning programs during development and regeneration, but with distinct initiation mechanisms

In initiating cell type specification programs, stem cells likely make decisions in the context of spatial patterning information. A gene regulatory network for posterior patterning has been probed functionally during regeneration in *H. miamia*, and we sought to study it during development. Specifically, amputation results in the wound-induced, polarized expression of genes (*wnt-1*, *wnt-3*, and *sp5*) at the posterior facing wound site (which will form the new tail). These genes are required for the specification and maintenance of posterior identity until a new tail and homeostatic patterning gradients are restored^27,32^ (Figure 4A). Given the similarity of temporal expression between development and regeneration in the bulk RNA-seq data (Figure 1G), we investigated the role of this network in anterior-posterior axis patterning during embryogenesis.

**Figure 4.**
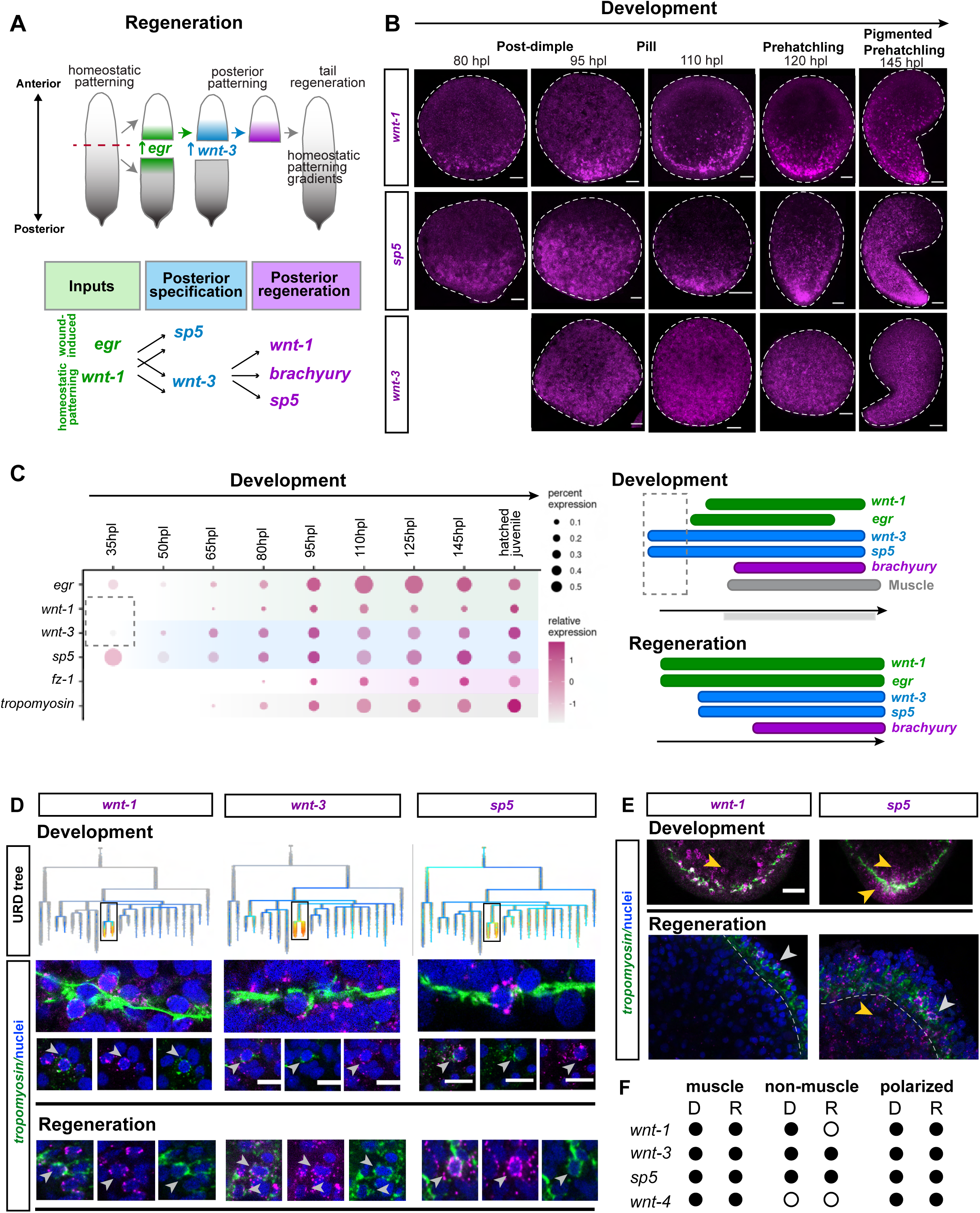
Spatial and cell-type expression suggest shared posterior patterning programs during development and regeneration, but with distinct initiation mechanisms. A) Summary of regenerative posterior patterning program from Ramirez et al 2020^27^. Following amputation, *egr* is wound-induced at both wound sites (green). In the posterior facing wound site, *egr* expression is required for *wnt-3* expression (blue), as is homeostatic *wnt-1.* Wound-induced *wnt-3* is required for expression of downstream posterior patterning genes (purple), restoration of homeostatic patterning gradients, and regeneration of the tail. B) *wnt-1, sp5,* and *wnt-3* are polarized during development from the post-dimple stage through pigmented prehatchling. C) Dot plot demonstrating expression of posterior patterning genes during development. During development, although all genes show robust expression around 80 hpl (late post-dimple), *wnt-3* and *sp5* are expressed prior to *egr* and *wnt-1*, which are required for *wnt-3* and *sp5* wound-induced expression during regeneration. D) Expression of *wnt-1, wnt-3,* and *sp5* on the URD tree shows high expression during development in muscle lineages (boxed). Posterior patterning genes (purple) are co-expressed with the pan-muscle gene *tropomyosin* (green) in both development and regeneration. E) During development, *wnt-1* is expressed in muscle regions (green – *tropomyosin*) as well as internal cell populations. During regeneration, it is restricted to the muscle regions. *sp5* is expressed in muscle and non-muscle regions of the animal in both development and regeneration. Dashed line shows boundary of muscle layer in regeneration images. F) Summary of features of developmental and regenerative expression of posterior patterning genes. Black circle indicates feature is present; white circle indicates not detected.

First, we examined the spatial distribution of these genes during development and found that they were polarized on one side of the embryo starting at the post-dimple stage (80 hpl) (Figure 4B). As the embryo elongates and anterior structures such as the pharynx and mouth become visible, this polarized expression is present at the opposing pole in posterior regions of the animal (Supplemental Figure 4A). This spatial expression, combined with the highly conserved role of Wnts in axial patterning across metazoan taxa^33,34^, supports a conserved role of these genes in *H. miamia* posterior patterning during development, as they serve during regeneration^27^.

We next examined the possibility that these genes are being deployed as part of the same GRN as during regeneration. Bulk transcriptome data suggested genes involved in this network became expressed ∼80 hpl in the post-dimple stage, but scRNA-seq data showed substantial *wnt-3* and *sp5* expression prior to this, which may have nonetheless been too low to detect with bulk sequencing (although *wnt-1* only became detectable close to 80 hpl in both data types) (Figure 4C). This temporal order suggests *wnt-1* is downstream of *wnt-3*, which contrasts with the regenerative GRN, where homeostatic *wnt-1* expression is required for initiating the expression of *wnt-3*. Meanwhile, *egr,* which is required for the expression of *wnt-3* and *sp5* during regeneration, does not exhibit high expression until after *wnt-3* and *sp5* are expressed, suggesting it is not needed for initiating their expression in development. Overall, this analysis suggested that *wnt-1*, *wnt-3*, and *sp5* are likely patterning the posterior in embryos, but the regulatory inputs controlling their expression may be different.

Given these temporal differences between development and regeneration, we next used scRNA-seq data to compare the cellular contexts of these genes in both processes. Previous work identified muscle as an important patterning tissue in *H. miamia* adults that expresses components of the posterior patterning network^27,35^. Double FISH of posterior patterning genes in conjunction with the muscle marker *tropomyosin* showed that the components of this network (*wnt-1, wnt-3*, and *sp5*) were expressed within muscle during regeneration (Figure 4D).

Extended to embryos, this approach showed that these genes were also expressed in muscle during development, which is corroborated by expression of these genes in the muscle lineages in the URD tree (Figure 4D). However, differences emerged when considering expression in cells other than the muscle cell population – *wnt-3* and *sp-5* were expressed in non-muscle cells in both development and regeneration, whereas *wnt-1*, exhibiting minimal expression outside of muscle during regeneration, showed much broader expression during development (Figure 4E, Supplemental Figure 4B). Notably, this expanded developmental expression was specific to *wnt-1* and not simply a feature of all posterior genes, as the marker *wnt-4*^36^ showed very clear restriction to within muscle cells in embryos (Supplemental Figure 4C). Given that the non-muscle *wnt-1+* cells during embryogenesis are expressed in internal regions where the stem cells are ultimately located^21,25^, it is possible that these are progenitor cells that will differentiate into muscle, ultimately affecting posterior patterning. Alternatively, it could be that additional cell types or progenitor cells express *wnt-1*, and are participating in mediating posterior patterning. Further, *wnt-1* expressed in non-muscle cells may not be involved in posterior patterning, although it also exhibits polarized expression in these non-muscle regions of the animal (Supplemental Figure 4B).

Overall, while our spatial studies of expression suggest that the posterior patterning program identified during regeneration is likely operational during development, our investigations of temporal expression dynamics and cellular contexts suggests that some of these genes may have broader roles in embryonic development and that they are regulated via distinct mechanisms.

### Chromatin profiling identifies development- and regeneration-specific regulatory DNA, but also reveals substantial utilization of shared regulatory elements for genes expressed in both processes

Given that our analyses indicated distinct initiation mechanisms and cell-type expression for both neural differentiation trajectories and posterior patterning during development and regeneration, we next sought to identify upstream genomic regulatory elements that could lead to these differences in gene expression. Chromatin accessibility can serve as an indicator of cis-regulatory function^37^, and our previous work utilizing ATAC-seq^38^ for chromatin profiling identified dynamic chromatin during regeneration in *H. miamia*^22^. Combining this assay with studies of gene function revealed gene regulatory networks for the wound response^22^, posterior patterning^27^, and neural specification^28^. Additionally, we generated chromatin profiling data from developing embryos, which revealed regulatory elements and transcription factor (TF) motifs associated with major processes during embryogenesis^26^. Leveraging these datasets, we sought to determine the extent to which development-specific, regeneration-specific, and shared regulatory elements drive the expression of the same genes during development and regeneration.

First, to broadly characterize the regulatory landscapes underlying development and regeneration, we queried the overlap between “peaks”, i.e. significantly accessible chromatin regions (Figure 5A). We focused this analysis on chromatin that was dynamic during regeneration as a proxy for regulatory regions that may be specifically associated with regenerative (as opposed to homeostatic) gene expression^22^, and on chromatin that was accessible at any sampled time point of development, reasoning that any of these peaks could be important for developmental gene expression. Genome-wide, we found a large number of peaks that were evenly spread across three categories: development-specific peaks, regeneration-specific peaks, and regions accessible during both processes (referred to onwards as “shared” peaks) (Figure 5B). We associated these peaks to the nearest gene loci using UROPA^39^, and found that slightly higher proportions of shared peaks were associated with protein-coding genes (Figure 5B). Of the 7,376 genes that had peaks assigned, around 30% had only regeneration-specific peaks whereas around 15% had only development-specific peaks (Figure 5C). We wondered whether shared peaks were primarily reflective of promoters being accessible, and found that while indeed promoter-associated peaks were a higher proportion of the shared peak set, this set also included regulatory regions associated with intronic and intergenic genomic features (Supplemental Figure 5A). Overall, the regeneration-specific peaks identified here represent putative regulatory mechanisms that operate uniquely relative to development, and future functional studies will establish the significance of their functions. Notably, binding site enrichment analysis of regeneration-specific, development-specific, and shared peaks did not uncover any striking differences in the types of transcription factors acting in these regulatory regions (Supplemental Table 3).

**Figure 5.**
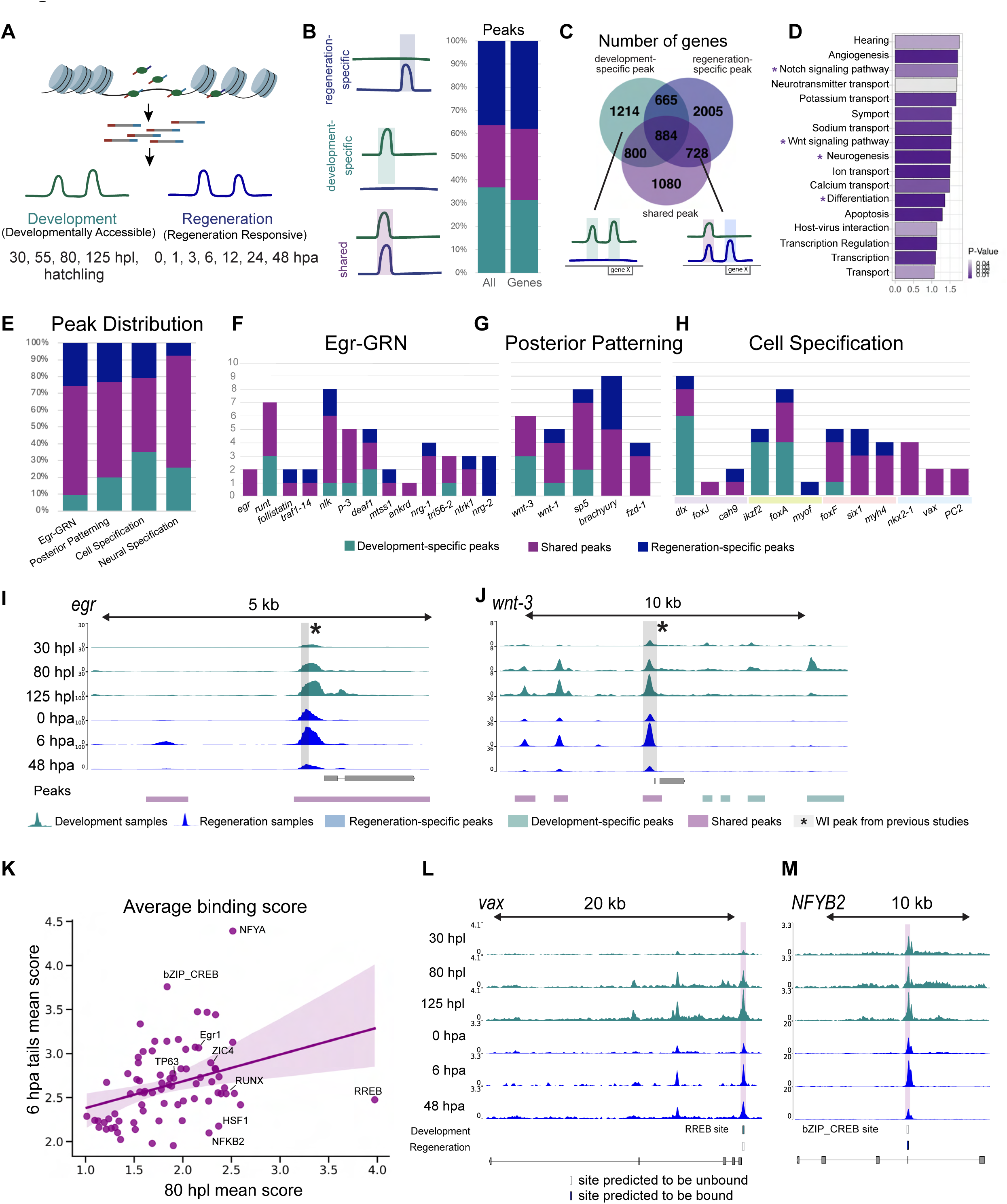
Chromatin profiling identifies development- and regeneration-specific regulatory DNA, but also reveals substantial utilization of shared regulatory elements for genes expressed in both processes. A) Previous studies produced both developmental and regenerative bulk ATAC-seq libraries in *H. miamia* that were utilized in this study. In brief, during ATAC-seq Tn5 transposase binds, cuts, and ligates sequencing adapters at locations in accessible DNA. These barcoded DNA fragments can be sequenced, aligned to the genome, and utilized to identify regions of accessible chromatin^38^. Any developmentally accessible region from 30 hpl, 55 hpl, 80 hpl, 125 hpl, or hatchling datasets was considered^26^. From the regeneration datasets for 0, 1, 3, 6, 12, 24 and 48 hpa in head and tail fragments we maintained peaks that showed changes in accessibility between two timepoints, in an attempt to distinguish between regenerative and homeostatic peaks^22^. B) Peaks from both datasets were identified as being called in only one dataset or only in either the regenerative or developmental peakset (left: blue - only in regeneration; teal - only in development, purple - both). Regeneration-specific (blue), development-specific (teal), or shared (purple) peaks were identified for all peaks (middle) or peaks associated with genes (right). C) Distribution of types of peaks (regeneration-specific, development-specific, or shared) was assessed for all genes with peaks associated with them. The number of genes that had only development-specific peaks, only regeneration-specific, only shared, or some combination of these categories was counted. D) GO terms for any gene associated with a peak present in both development and regeneration. Asterisks represent terms associated with processes shared between development and regeneration that have been studied in *H. miamia*. E) Distribution of types of peaks (development-specific, regeneration-specific, or shared) for genes in the Posterior patterning network, Cell specification, Neural specification, or Egr-GRN. F-H) Distribution of types of peaks (development-only, regeneration-only, or shared) for each gene in the Egr-GRN (F), Posterior patterning network (G), or Cell specification (H). In (H) purple marks epidermal genes, green are endodermal genes, salmon are muscle genes, and light blue are neural genes. I-J) Chromatin accessibility of the *egr* (I) or *wnt-3* (J) locus during development (teal) and regeneration (blue). Peaks that were only detected in the development dataset (teal), regeneration dataset (blue) or were detected in both (purple) are shown. *egr-r*esponsive peak from regeneration^22^ is highlighted in gray with an asterisk. K) Linear regression plot comparing mean binding scores between 6 hpa tails in regeneration and 80 hpl embryos during development. Motifs with high deviation from diagonal are highlighted (RREB, HSF1, NFKB, bZIP_CREB), as are previously investigated motifs (Egr, RUNX, TP63, ZIC4)^26,22^. Shaded blue line: 95% confidence interval. Slope = 0.304, y-intercept = 2.07, R^2^ value = 0.337, p-value = 0.00257. L-M) Chromatin accessibility of the *vax* (L) or *NFYB* (M) locus during development (teal) and regeneration (blue). Location of motif binding sites for RREB (L), *vax)* or bZIP_CREB (M), *NFYB*) are highlighted in purple and marked with rectangles. Sites predicted to be bound in a process are shown with a filled in rectangle, sites predicted to be unbound in a process are indicated with an empty rectangle.

Recognizing that the numbers of development- or regeneration-specific peaks detected is highly dependent on extent of sampling, e.g. seven stages over 48 hours of regeneration were sampled in contrast to five stages over ∼175-190 hours of development (Figure 1A), we next focused our attention on the shared peaks. Genes containing both regeneration-responsive and developmentally-accessible chromatin were enriched for functions in many development-associated processes such as Notch signaling, Wnt signaling, and differentiation as well as in cellular processes such as neurotransmitter transport and apoptosis (Figure 5D, Supplemental Table 4). We next focused on the subset of these genes whose functions had been previously characterized in regenerating worms, examining the loci of genes in the posterior patterning, cell-type specification, and Egr-GRN networks to understand their regulation. Although these genes contained all three categories of regulatory regions, the proportion of regeneration-specific peaks was substantially lower relative to genome-wide levels (Figure 5E-H, Supplemental table 5). Strikingly, we found that a number of these genes contained no regeneration-specific peaks, including several well-characterized wound-induced genes (such as *egr* and *runt*) (Figure 5F, I-J). In particular, the previously described regenerative-responsive peaks containing Egr motifs also displayed accessibility during development (Figure 5I, J, Supplemental Figure 5B). Consistent with similarities we had observed in cell specification, we observed a number of shared peaks at the loci of genes in these processes between development and regeneration (Supplemental Figure 5B). Altogether, the finding that substantial proportions of regeneration-responsive chromatin are also accessible during development genome-wide, including the previously-studied regeneration-responsive regions in wound-induced genes, suggests that shared regulatory regions play important roles in driving regenerative expression of developmental genes.

Given the presence of shared regulatory regions, we next asked how different inputs, as regeneration and development must have, achieve activation of gene expression during the two processes. To address this question, we asked whether similar or different TF motifs could be playing a role within shared regulatory regions during development and regeneration. Each accessible chromatin region typically contains multiple TF binding motifs, which in some cases overlap with each other. It is noteworthy, however, that the presence of a motif alone does not indicate when, or if, the corresponding TF may actually bind there and be involved in modulating transcription at that locus. Using TOBIAS^40^, we predicted TF binding via footprinting analysis of binding sites within shared peaks. Pairwise comparisons of predicted TF binding scores between two developmental or two regenerative timepoints are tightly correlated, with R^2^ values >0.9 (Supplemental Figure 5C, Supplemental Table 6). In contrast, we found a more modest correlation of mean binding scores between motifs in developmental and regenerative timepoints (with R^2^ values between .3 and .6) (Fig. 5K, Supplemental Figure 5C, Supplemental Table 6). This correlation is driven by a number of motifs that appear to have similar relative mean binding scores between development and regeneration, which is to be expected. For example, our work shows that cell specification trajectories are shared and driven by the same TFs in both development and regeneration, and in fact we found that in some cases there is predicted binding of these TFs in shared regulatory regions (Supplemental Figure 5D).

However, some motifs are “off-diagonal”, indicating that binding is preferentially high in one process and relatively lower in the other (Figure 5K). For example, RREB motifs are more highly bound in development whereas bZip_CREB motifs are more highly bound during regeneration genome-wide. In concordance with this, we find loci with differential predicted binding of these motifs within a shared peak between development and regeneration. For example, *vax*, which likely plays a similar role in neural specification in *H. miamia* development and regeneration but exhibits broader spatial expression early in development^28^ (Supplemental Figure 3B), is predicted to be bound by RREB in development but not regeneration (Figure 5L). On the other hand, the wound-induced gene *NFYB2*^28^ has sites predicted to be bound by bZIP_CREB during regeneration but not development, potentially explaining its regenerative activation (Figure 5M). Overall, these data suggest that even within shared regulatory regions, different TFs could mediate gene expression across development and regeneration.

## DISCUSSION

The question of how regeneration and development relate to each other has been long-standing in the field of developmental biology. Prior work focusing on specific experimental studies of vertebrate appendages have established that broad similarity can be accompanied by nuanced differences^12,13^. This work leaves open the question how broadly generalizable these findings are to other structures and to other organisms. Transcriptome-wide studies of regeneration in invertebrates have shown broad re-expression of developmental genes during regeneration, some putatively in the same pathways and others likely in different networks^9–11^. However, these studies don’t enable an explicit assessment of whether redeployed genes are performing the same functions and of the mechanisms that enable the re-expression of development genes. By applying single-cell transcriptome analysis and functional genomics in *Hofstenia miamia*, a species that is amenable to studies of both development and regeneration, we show that many genes expressed during regeneration are replaying their developmental functions. Further, we show that the genes performing functions in new pathways during regeneration are most likely still activated using the same regulatory regions that drove their expression during development (Figure 6).

**Figure 6.**
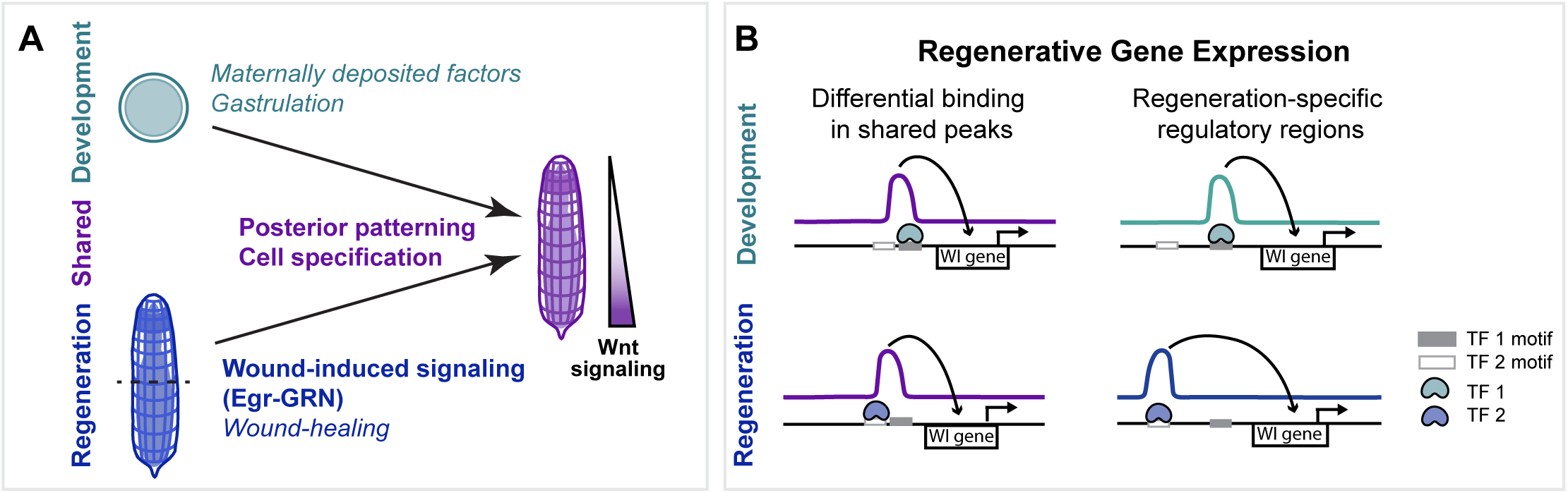
A) Despite different starting points, development and regeneration utilize similar pathways in *H. miamia* for cell specification and posterior patterning. A wound-induced network (the Egr-GRN) appears to have network architecture specific to regeneration. B) We propose that regeneration-specific expression is mediated by differential TF binding within regulatory regions shared between development and regeneration (left) as well as via distinct regulatory elements (right).

Our comparisons of transcriptome data in bulk recapitulated the results of previous similar studies in other systems, identifying many cohorts of genes that show temporal co-expression during both development and regeneration. However, to delve deeper into biological processes reflected in these results, we focused on genetic pathways that had been studied functionally during regeneration in *H. miamia*. Whereas posterior patterning genes showed correlated expression across both processes, wound response genes and cell type specification genes did not. Layering in single-cell transcriptome data, with accompanying experimental corroboration of developmental gene expression, showed that cell type specification programs are largely the same during development and regeneration, and the lack of correspondence observed in the bulk transcriptomic comparisons is likely driven by sampling. Thus the data points selected for bulk sequencing can significantly impact conclusions from comparisons of development and regeneration. We therefore suggest that some corroboration of gene expression and function are essential for rigorous statements in such studies.

Our analyses identified the Egr-controlled wound response gene regulatory network (Egr-GRN) as a possible novelty of regeneration, as we did not see consistent temporal or cell type co-expression of these genes during development, although these genes are expressed in both processes. This raised the question of how re-expression of genes during regeneration is mediated. Previous studies have identified regeneration-responsive regulatory elements that contribute to regenerative gene expression, however in most cases it is not clear which of these elements are regeneration-specific^4,41–47^. Our comparative analysis of regulatory landscapes revealed development- and regeneration-specific regulatory biology as well as substantial utilization of the same regulatory regions during both processes, including for genes in the Egr-GRN, where regenerative *egr*-responsive peaks had been identified and were found to be shared with development. Yet, different inputs must be needed to activate these genes during regeneration relative to development, and we found evidence suggesting that different binding sites within the same regulatory regions may be utilized predominantly in one process versus the other. This is reminiscent of a finding in *Drosophila* that different elements in the same enhancer were responsible for damage-responsive versus developmental expression of *wingless* in imaginal discs^4^. Functional studies that specifically perturb these binding sites will be needed to validate this hypothesis, and can be conducted in *H. miamia* as the species is amenable to CRISPR-Cas9 genome editing^48^.

We also identified substantial numbers of regeneration-specific regulatory regions, including in regions associated with genes that are also expressed during development. Together with the uniquely regeneration-responsive binding sites, these represent mechanisms that initiate expression of developmental programs during regeneration (Figure 6B). These mechanisms could serve as the ideal substrates for studies seeking to understand the evolution of regeneration mechanisms – generating similar data from other species, as others have done recently^15^, would enable a comparison of regeneration-specific mechanisms as opposed to comparing redeployed developmental pathways^13,19^. Future work could also reveal how generalizable our findings of the correspondence between development and regeneration in *H. miamia* are. In particular, studies in other species could assess whether the similarity of cell specification programs in this system, which uses a stem cell based differentiation process to make new tissue, extend to other species that utilize dedifferentiation-based programs.

## ACKNOWLEDGEMENTS

Many thanks to former and present members of the Srivastava and Koenig labs for insightful discussions regarding the work in this study for the duration of the project. A special thanks to Ryan Hulett, Paul Bump, Catriona Breen, Ye Duan and Carlos Rivera-Lopez for critical comments on the manuscript. This work was supported by the Harvard Bauer Core, FAS informatics and Harvard Research Computing. M.S. and K. L-S. were supported by the National Institutes of Health (R35GM128817, R35GM153252, and T32GM135143) and the Smith Family Foundation.

## AUTHOR CONTRIBUTIONS

The experiments were conceived and designed by K.L-S. and M.S. Experiments and analyses were performed by K.L-S. Figures were prepared by K.L-S. with input from M.S. The manuscript was written by K.L-S. and M.S.

## METHODS

### Data availability

Original data for RNA-seq analysis and ATAC-seq datasets in this study are available from multiple sources:

Regeneration bulk RNA-seq and ATAC-seq datasets^22^: NCBI BioProject PRJNA512373 Regeneration single cell RNA-seq^23^: NCBI Sequence Read Archive: PRJNA908236 Development bulk RNA-seq^24^: RNA-seq data are available at http://n2t.net/ark:/84478/d/fzht4sx3. Raw fastq files: NCBI Sequence Read Archive PRJNA603318.

Developmental scRNA-seq^25^: NCBI Sequence Read Archive PRJNA888438 Development bulk ATAC-seq^26^: upon request (will be deposited to NCBI SRA)

## EXPERIMENTAL MODEL AND SUBJECT DETAILS

The animals used in this study are the result of random matings from 120 adult *Hofstenia miamia* individuals collected in Bermuda in 2010^21^. Adult animals were cultured at 21° C in ∼1L of artificial seawater water (ASW) (37 parts per thousand of Instant Ocean Sea salt or reef crystals; ph 7.8-8.2) in plastic tupperware boxes with 20-40 adults per box. Embryos used in this study were collected from boxes, staged, and reared at 23° C in ASW. Twice a week boxes were cleaned, replaced with fresh ASW, and fed freshly hatched brine shrimp.

### Bulk RNA-seq analyses

Normalized TPM matrices of bulk RNA-seq datasets produced for Gehrke et al. 2019^22^ and Kimura et al. 2021^24^ were used for these analyses. Gene Ontology (GO) Analyses were performed using the DAVID tool^49^ as in Kimura et al. 2021^24^ using a custom *H. miamia* background set of genes and script to convert *H. miamia* transcript IDs to Human Uniprot IDs.

#### Fuzzy c-means clustering

A count matrix of normalized TPM values was used as input into the R program Mfuzz^50^ to group genes based on expression dynamics. As per the standard pipeline, null values were filtered and fuzzy-c means soft clustering was performed using fuzzifier *m* values and number of clusters that minimized overlap between clusters. Good cluster resolution was obtained in both development and regeneration using the default value of *m* = 1.25, with 10 clusters for development and 7 for regeneration.

#### Gene Overlap

Clustering assignments from fuzzy c-means clustering were used to extract lists of genes corresponding to each cluster. These lists were used as input into the R program Gene Overlap^51^ (version 1.38.0). Basic pipeline was used to identify sets of overlapping genes and significance of overlap for all pairwise comparisons. The R program circlize (version 0.4.15)^52^ was used to visualize shared gene sets between developmental and regenerative clusters using gene overlap matrix outputs from the Gene overlap program as input adjacency matrix.

### scRNA-seq analyses

#### Pearson Correlation

Pearson correlation was run as in Hulett et al. 2023^23^. Pseudobulk matrices were either produced by extracting count data from URD tree lineages from Kimura et al. 2022^25^ or from regeneration clusters as identified in Hulett et al. 2023^23^.

#### Lineage trajectory inference analyses

The *H. miamia* developmental URD tree from Kimura et al. 2022^25^ was used to visualize gene expression. To create the subsetted developmental neural tree we extracted count matrices and metadata from the neural URD branches once they split. These data were then clustered using the Seurat package^53^. The earliest time point was specified as the root and hatchling juvenile as the tip. For the regeneration dataset, neural clusters and neoblast clusters identified as being neural progenitors were subsetted and clustered. The H3.3+ neoblast cluster was used as the root and differentiated neural clusters (based on markers and analyses in Hulett et al. 2023^23^) as the tips. URD pipeline from Farrell et al. 2018^29^ was used to create the new URD tree. Spline plots were produced as in Primack et al. 2023^54^.

### FISH

Whole worms and regenerating fragments were fixed for 1 h in 4% PFA in PBST, on nutator, at room temperature. Embryos were dechorionated as in (Kimura, et al. 2021)^24^ (32 mM sodium hydroxide, 0.5 mg/ml sodium thioglycolate and 1 mg/ml of pronase in artificial seawater (ASW) then fixed in 4% PFA in ASW, on nutator, at room temperature. Specimens were washed in PBST and either kept at 4°C for up to one week (embryos) or stored in 100% methanol at -20° C until use (whole worms and regenerating fragments).

Fluorescence *in situ* hybridization (FISH) was performed following the protocol in Srivastava et,al 2014^21^ (whole worms and regenerating fragments) or Kimura, et al. 2021^24^ (embryos). Digoxigenin and Fluorescein labeled riboprobes were prepared as in Srivastava, et al. 2014^21^. Minor modifications from Srivastava 2014 were: 1) All washes were performed with 800uL of solution; 2) Bleach solution was changed to incubation in solution (5% deionized formamide,1.2% H2O2, 50% saline-sodium citrate buffer (SSC) in milli-Q water) under a strong light source for 2 hours, 3) Prior to antibody treatment animals were incubated in block solution for 1-2 h. Nuclear staining was performed by either incubating in DAPI® (VWR, 80051-386) (1 μg ml^−1^) for 45 minutes (hatched worms; embryos in which AlexFluor 647 secondary antibody was used) or for 45 minutes in TO-PRO-3 Iodide (642/661) (Thermo Scientific) (embryos). Specimens were mounted in VECTASHIELD® (VWR, 101098-042). Imaging was performed using a Leica SP8 confocal microscope.

### Immunohistochemistry

Prior to fixation, embryos were dechorionated as outlined in Kimura et al. 2021^24^ and above. Whole worms, regenerating fragments, and embryos were fixed in 4% paraformaldehyde in ASW rocking for 1 hr at room temperature then moved to PBST. Animals were incubated for 30 min in block solution (10% horse serum in PBST), then incubated in primary antibody diluted in block solution for 48 hours at 4°C (FMRFamide: EMD Millipore AB15348, 1:300; custom *H.miamia* Tropomyosin antibodies^30^ used together at a final concentration of 1 μg ml^−1^)).

Specimens were washed 8×20 minutes in PBST at room temperature, then incubated overnight at 4°C in secondary antibodies diluted 1:200 in block solution (AlexaFluor488 goat anti-rabbit IgG (Jackson Immunoresearch 111-545-144); AlexaFluor568 goat anti-rabbit (Abcam 175471); or AlexaFluor647 (Jackson ImmunoResearch Laboratories, 111-545-003)). Specimens were washed 8×20 minutes in PBST. Nuclear staining was performed by either incubating DAPI® (VWR, 80051-386) (1 μg ml^−1^) for 45 minutes (hatched worms; embryos in which AlexFluor 647 secondary antibody was used) or for 45 minutes in TO-PRO-3 Iodide (642/661) (Thermo Scientific). Specimens were mounted in VECTASHIELD® (VWR, 101098-042).

### ATAC-seq Analysis

#### Identification of features shared between and unique to each dataset

Consensus peaksets for developmentally dynamic peaks (Bump & Loubet-Senear, at al.^26^) and regeneration-responsive peaks (Gehrke et al, 2019^22^) were used to identify peaks present in the developmental dataset, the regenerative dataset, or both. To identify shared regions, the BEDtools^55^ command intersectBed was used with the arguments -wa -a to retain shared features, and the arguments -v -a to find features present in only one dataset.

#### Downstream peakset comparisons

UROPA^39^ was used to link peaks to the neighboring gene using default parameters. The cutoff to link a peak to an annotated feature in the genome was 10kb. Resulting lists (development-only, regeneration-only, and shared) were used as input into the R package VennDiagram^56^ (version 1.7.3) to visualize overlap and extract sets of peaks from each category. GO analyses were performed using the DAVID tool^49^ as described in Kimura et al. 2021^24^, using as input lists of transcripts linked to peaks using UROPA.

ChIPseeker^57^ was used to identify distribution of peaks (promoter, intron, exon, intergenic), using as inputs the development-only, regeneration-only, or shared peaksets. The analysis was performed using either all peaks in each of these categories or only peaks linked to genes using UROPA. Predictions of motifs enriched in sets of peaks were done using the SEA^58^ tool on MEME suite^59^. The BEDtools^55^ command getfasta was used to convert .BED files corresponding to each set of peak regions to FASTA format to use as input into SEA. A curated *H.miamia* motif list^28^ was used as input for motifs.

#### Binding scores of shared peaks

Merged bam files from Bump & Loubet-Senear et al.^26^ or Gehrke et al.^22^ were fed into the TOBIAS pipeline and analyzed using the standard pipeline as outlined in Bentsen et al.^40^, with minor modifications as in Bump & Loubet-Senear et al.^26^. The input peakfile was the “shared” peakset. In short, we ran TOBIAS ATACorrect, FootprintScores, and BINDetect. Mean binding scores were found by comparing all head, all tail, or all development samples using the BINDetect function. The python program Seaborn^60^ was used to calculate and plot correlation between mean binding scores using the regplot() function. ATAC-seq tracks were visualized using pyGenomeTracks^61,62^

## SUPPLEMENTAL FIGURE LEGENDS

**Supplemental Figure 1.** A) Jaccard Index and number of genes from each pairwise comparison of fuzzy c-means between development and regeneration clusters. B) Overlap between cluster R1 and developmental clusters. i) Chord plot highlighting contribution of genes from developmental clusters to regenerative cluster R1. Developmental clusters marked in teal were identified as having greater overlap than by random chance. ii-vi) GO terms associated with selected overlapping clusters (highlighted in teal on the chord plot). Fuzzy cluster profiles for developmental and regenerative clusters being compared are shown above the GO term plots. C) Heatmaps of regenerating tails for genes in candidate networks.

**Supplemental Figure 2.** A) Expression of genes on the developmental URD tree for additional tissue markers that were identified and functionally interrogated during regeneration in Hulett et al. 2023^23^.

**Supplemental Figure 3.** A) At hatching (left), *H. miamia* exhibits many features of the nervous system described in the adult (right), including an anterior condensation of neurons, a subepidermal nerve net, and a dorsal commissure (shown by *gad-1*), and clumps of nuclei that are between neurite bundles. B) Developmental expression of additional neural markers via FISH and on the developmental URD tree. C) Immunohistochemistry for FMRFamide showing the nerve net during development, first at 145 hpl where it seems sparse, and later at 170 hpl where the distribution resembles that in hatched juvenile worms. D) *soxC/pou4* and *pou4/pkd2* are co-expressed and enriched in the same URD branches as each other in development and regeneration. E) Whole mount images of *soxC/nkx2-1* and *nkx2-1/trpC-1* corresponding to 63X images in Figure 3. F) Expression of NFY family genes on the developmental URD tree. Expression is broad with varied cell type expression. G) Expression of Egr-GRN members on the developmental URD tree. Scale bars: 20x images, 10um; 63x images, 50 um.

**Supplemental Figure 4.** A) Wnts are polarized in posterior regions of the animal once morphological features such as the pharynx (ph, anterior) and pointy tail (posterior) are present. B) *wnt-1* is expressed in internal, non-muscle cells from at least the post-dimple stage. Images are single slices from a z-stack in the middle of the animal. C) At all time points surveyed, *wnt-4* is only expressed within muscle cells based on FISH (left) and expression in the developmental URD tree (right), and does not exhibit expression in internal cells as *wnt-1* does. White arrowheads, co-expression between *wnt-4* and the muscle marker *tropomyosin.* Scale bars: 20x images, 10um; 63x images, 50 um.

**Supplemental Figure 5.** A) Bar plots showing the proportions of peaks in the promoter, introns, intergenic regions, and exons for sets of peaks identified only in the regeneration peakset, development peakset, or present in both peaksets for (left) all peaks and (right) peaks associated with genes via UROPA. Far right: schematic depicting categories of peaks in bar plots. B) Chromatin accessibility of additional genes in posterior patterning, cell specification, neural differentiation, or Egr-GRN processes. Teal tracks: developmental timepoints, blue tracks: regenerative timepoints. Shaded gray regions are previously validated wound-induced peaks^22,27,28^ C) Linear regression plot for all pairwise comparisons of binding scores between libraries in developmental and regenerative ATAC-seq time courses. D) Top: Predicted co-expression in the *H.miamia* developmental URD tree between *foxA* and *fabp5,* and *foxF* and *myh4*. Boxed URD tree lineages correspond to known regenerative cell-type expression of these genes (*foxA/fabp5*: endodermal; *foxF/myh4:* muscle). Bottom: Chromatin accessibility of the *fabp5 and myh4* loci during development (teal) and regeneration (blue). There is a predicted Fox binding site in a shared peak of *fabp5*, and *myh4*, which in the future could be interrogated to assess direct binding of *foxA* or *foxF* at these loci in development and regeneration.

