## Supplemental Figures for "Regeneration recapitulates many embryonic processes, including reuse of developmental regulatory regions"

### Supplemental Figure 1

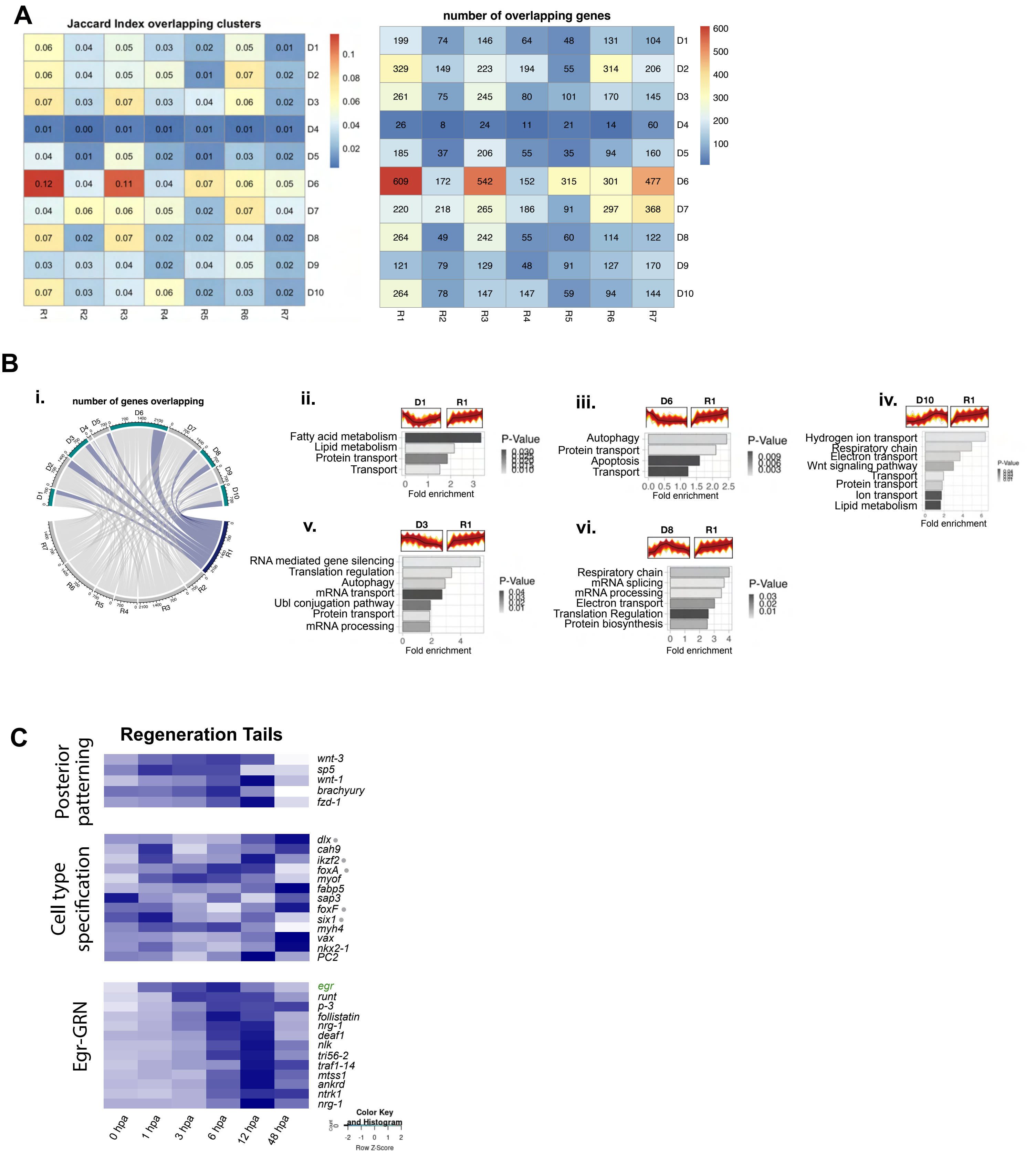

#### Supplemental Figure 2

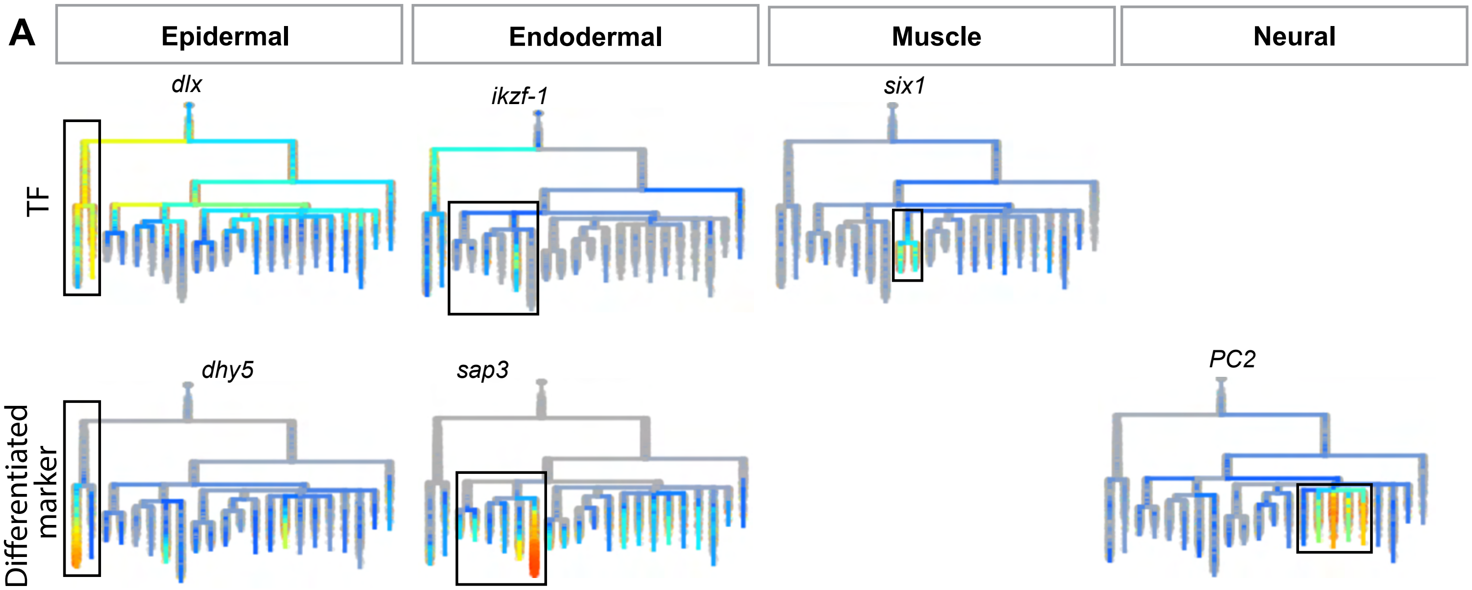

#### Supplemental Figure 3

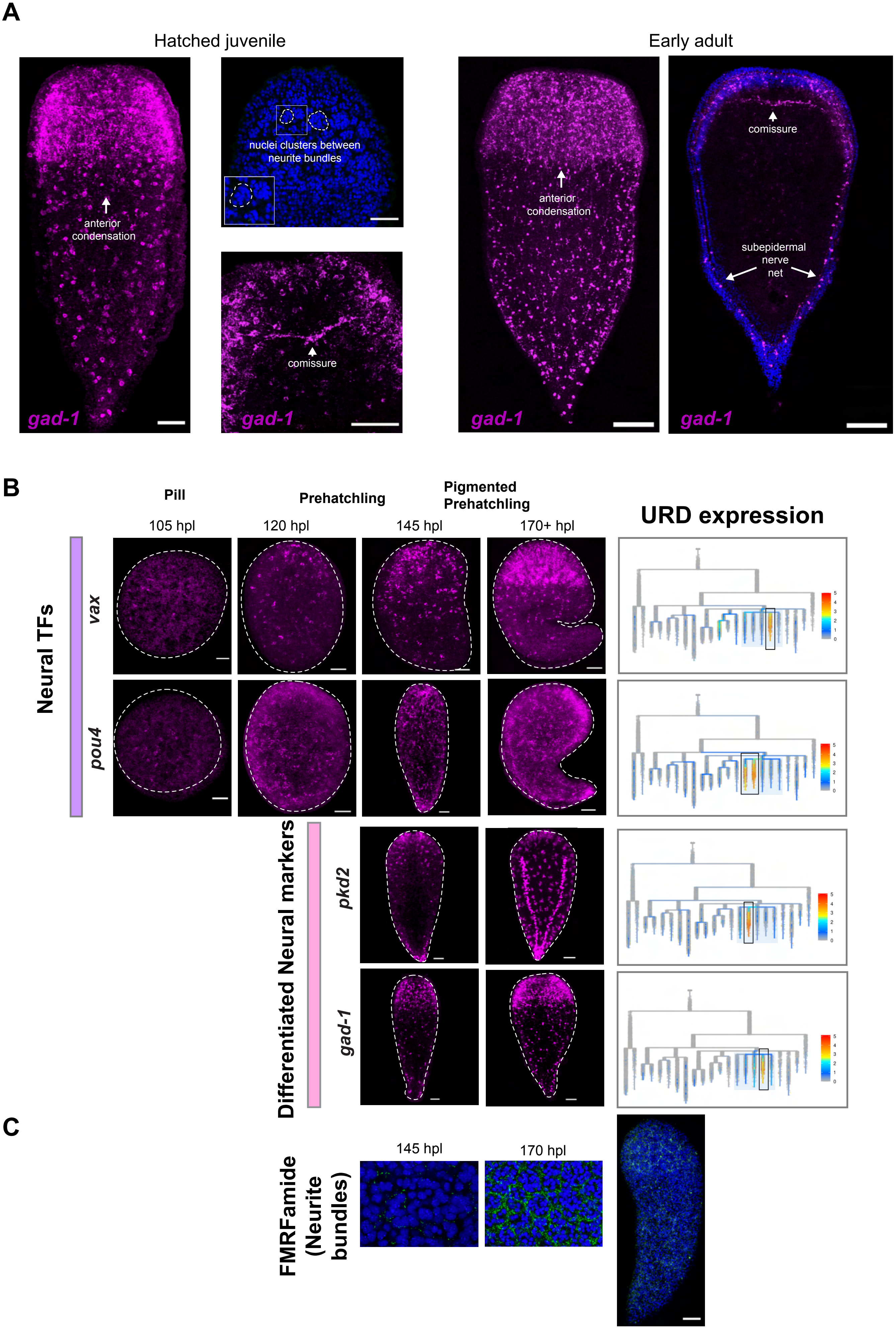

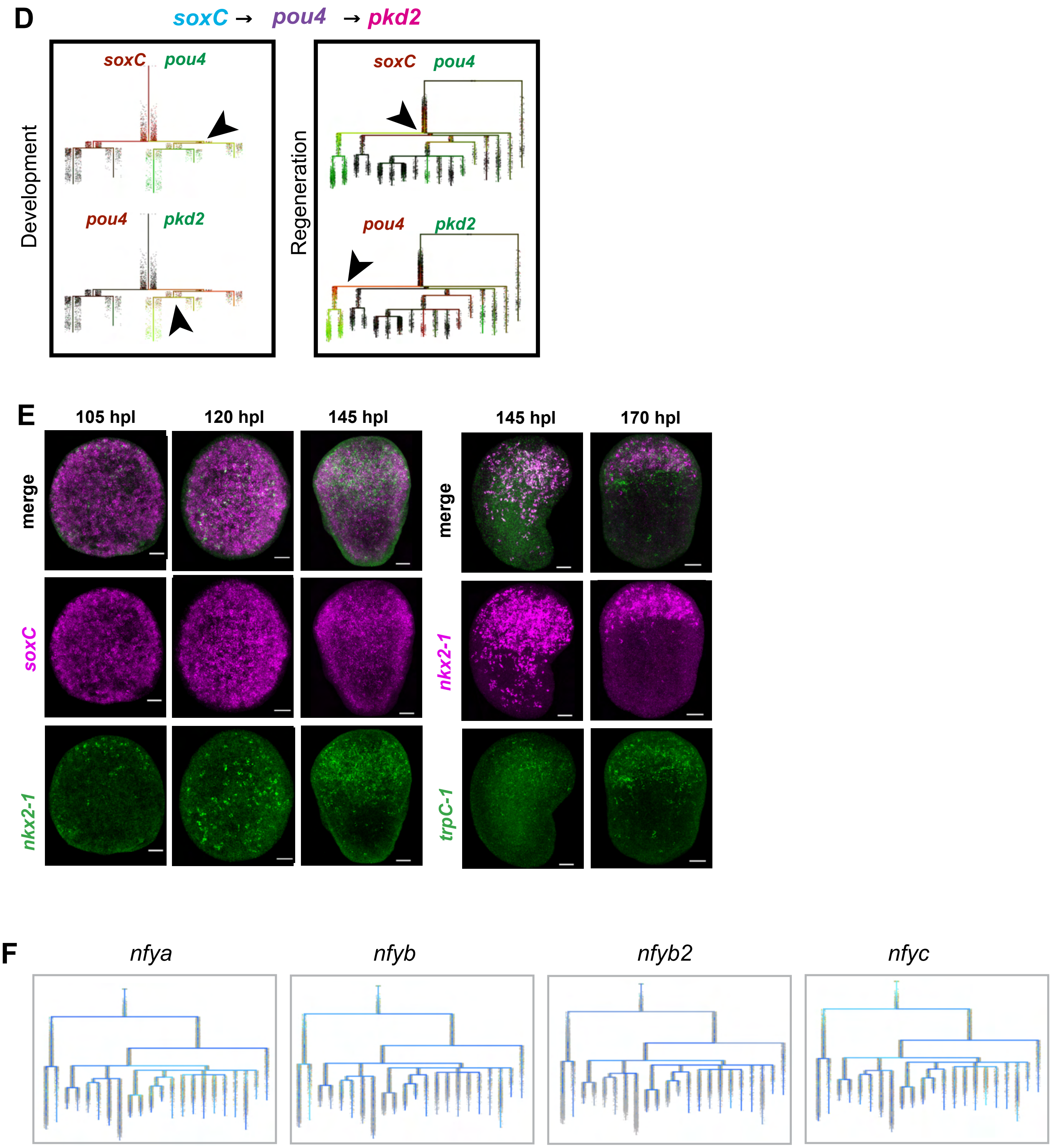
D
soxC → pou4 → pkd2
Development
soxC pou4
pou4 pkd2
Regeneration
soxC pou4
pou4 pkd2
E
105 hpl
120 hpl
145 hpl
145 hpl
170 hpl
merge
soxC
nkx2-1
merge
nkx2-1
trpC-1
F
nfya
nfyb
nfyb2
nfyc

G

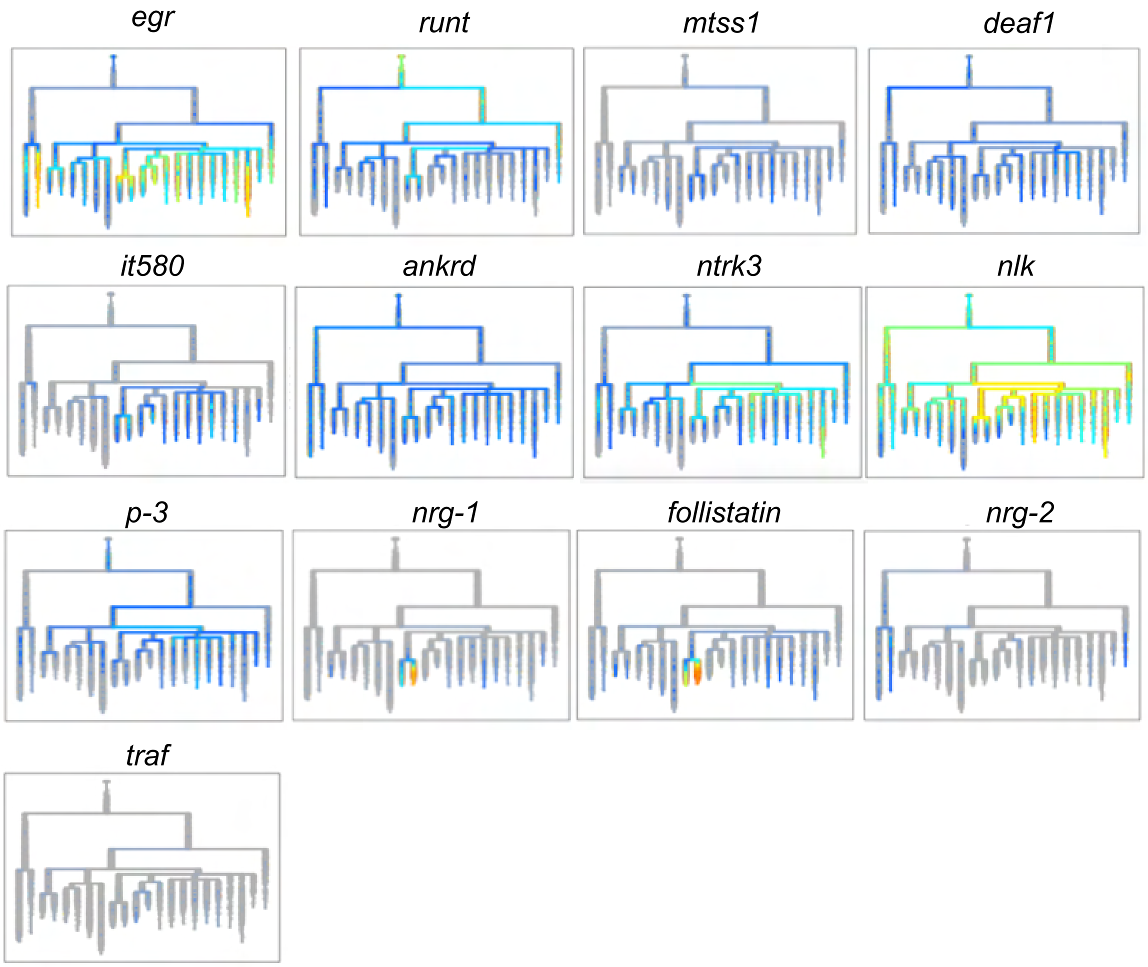

egr
runt
mtss1
deaf1
it580
ankrd
ntrk3
nlk
p-3
nrg-1
follistatin
nrg-2
traf

### Supplemental Figure 4

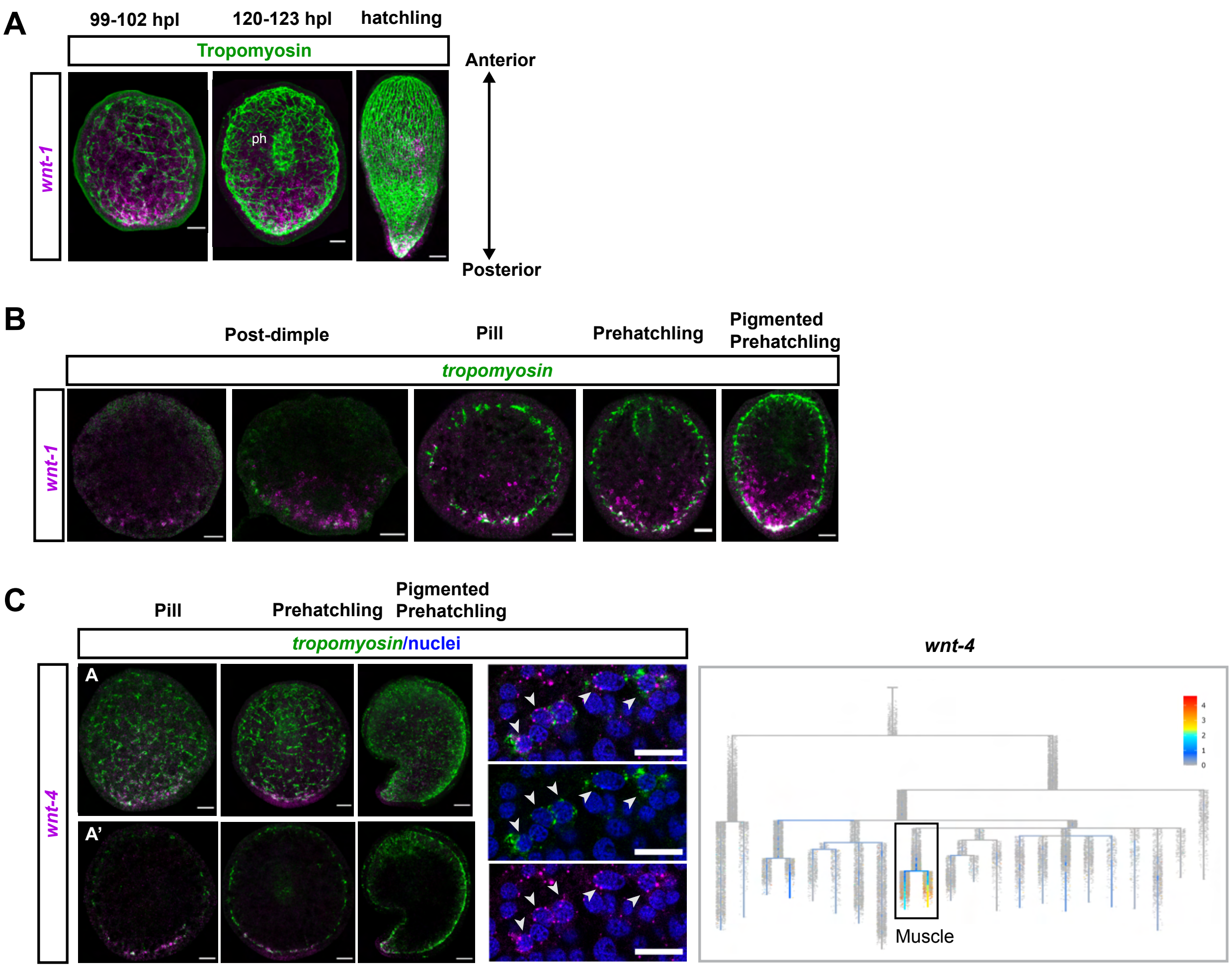

#### Supplemental Figure 5

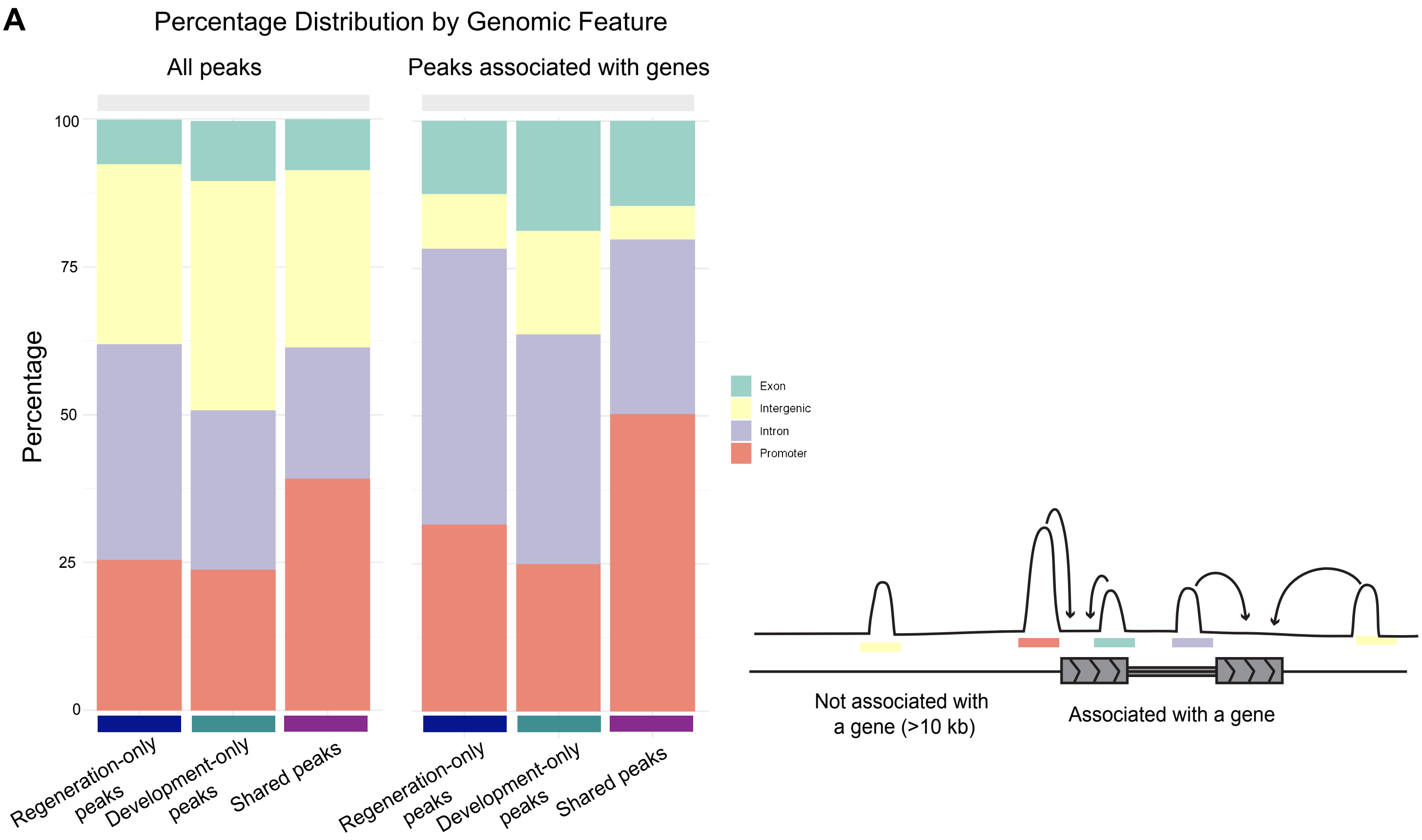

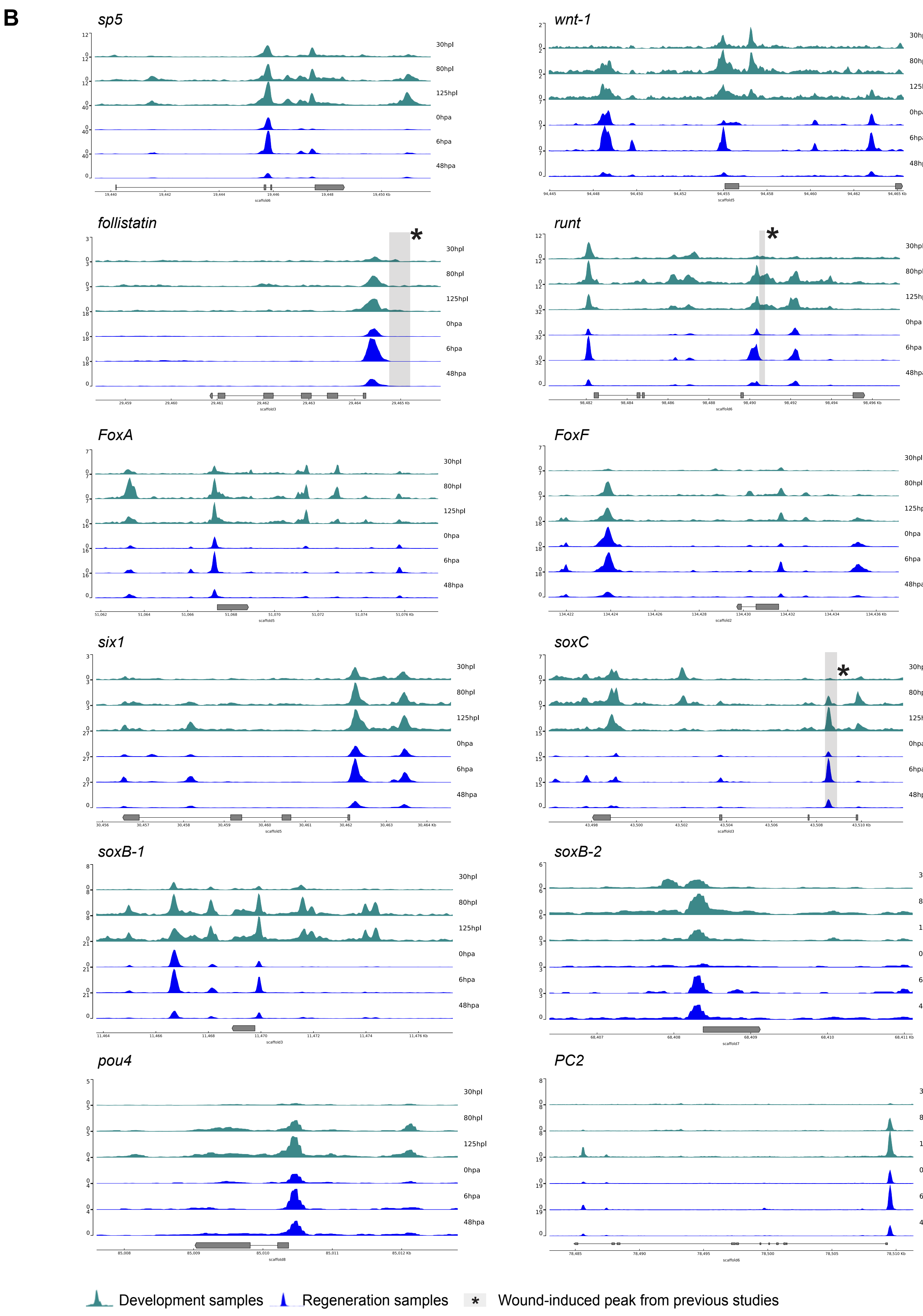
B
sp5
wnt-1
follistatin
runt
FoxA
FoxF
six1
soxC
soxB-1
soxB-2
pou4
PC2
30hpl
80hpl
125hpl
0hpa
6hpa
48hpa
*
Development samples
Regeneration samples
Wound-induced peak from previous studies

**C**

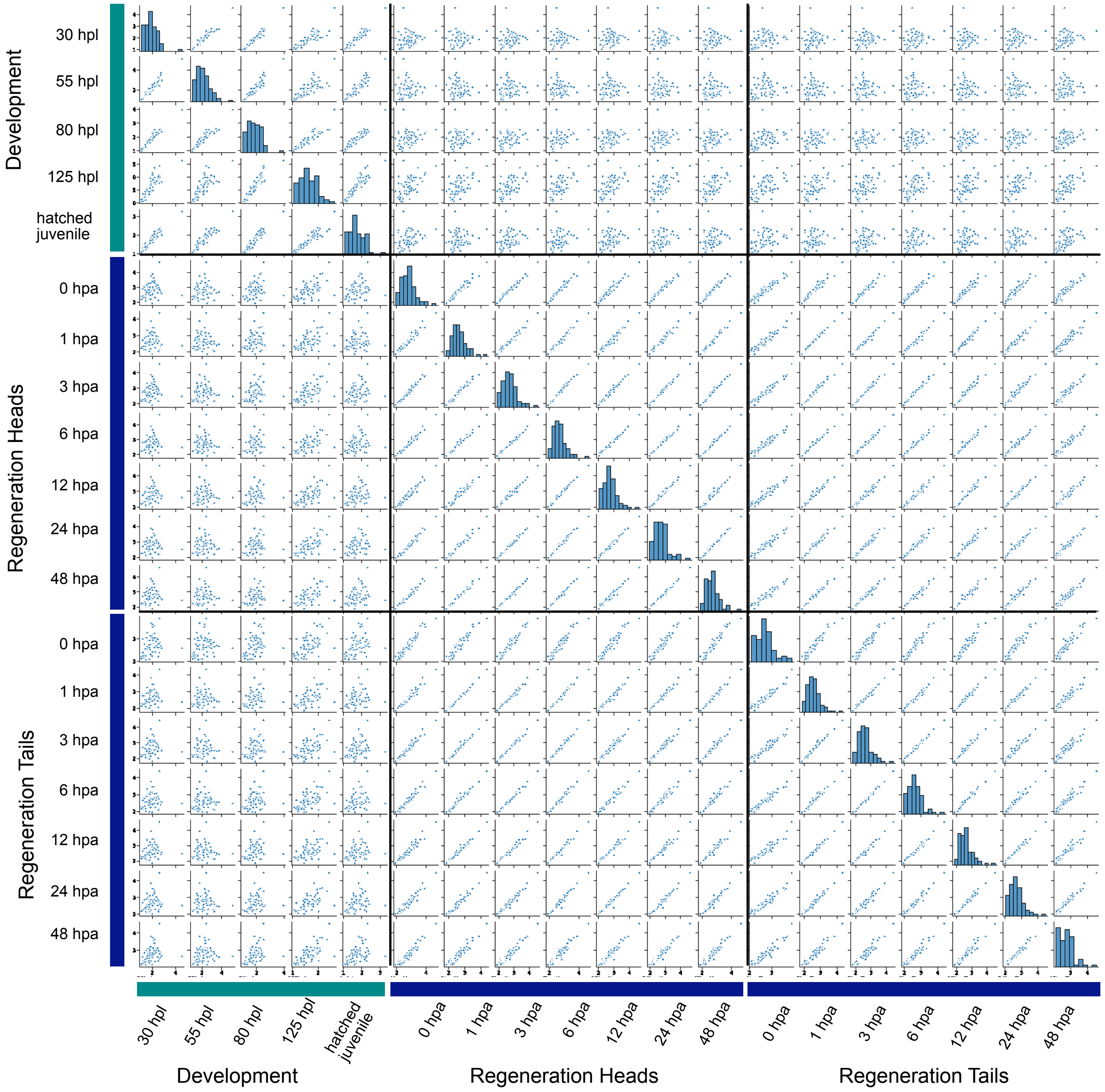

Development
30 hpl
55 hpl
80 hpl
125 hpl
hatched juvenile
Regeneration Heads
0 hpa
1 hpa
3 hpa
6 hpa
12 hpa
24 hpa
48 hpa
Regeneration Tails
0 hpa
1 hpa
3 hpa
6 hpa
12 hpa
24 hpa
48 hpa
30 hpl
55 hpl
80 hpl
125 hpl
hatched juvenile
0 hpa
1 hpa
3 hpa
6 hpa
12 hpa
24 hpa
48 hpa
0 hpa
1 hpa
3 hpa
6 hpa
12 hpa
24 hpa
48 hpa
Development
Regeneration Heads
Regeneration Tails

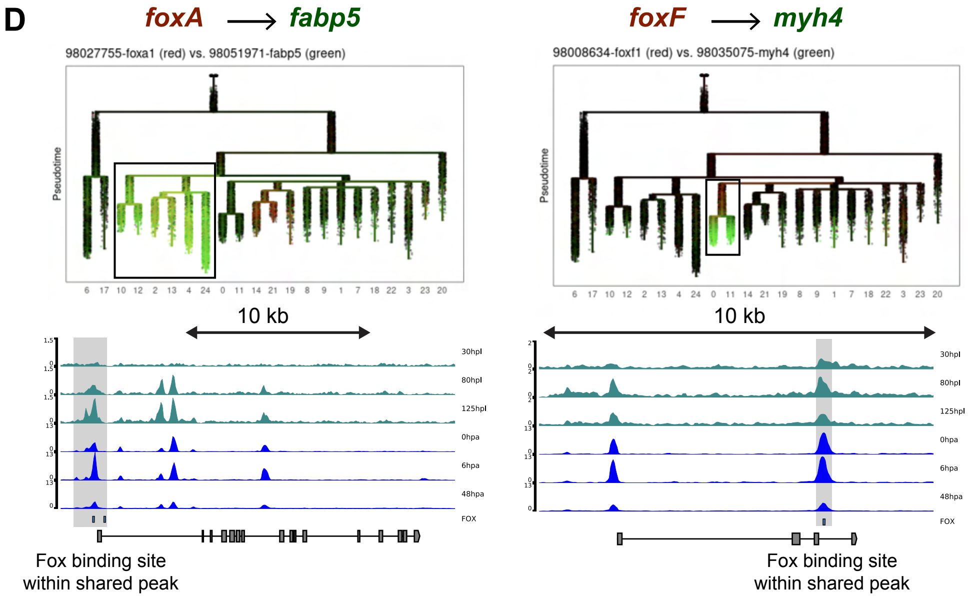
D
foxA → fabp5
98027755-foxa1 (red) vs. 98051971-fabp5 (green)
Pseudotime
6 17 10 12 2 13 4 24 0 11 14 21 19 8 9 1 7 18 22 3 23 20
10 kb
30hpl
80hpl
125hpl
0hpa
6hpa
48hpa
FOX
Fox binding site
within shared peak
foxF → myh4
98008634-foxf1 (red) vs. 98035075-myh4 (green)
Pseudotime
6 17 10 12 2 13 4 24 0 11 14 21 19 8 9 1 7 18 22 3 23 20
10 kb
30hpl
80hpl
125hpl
0hpa
6hpa
48hpa
FOX
Fox binding site
within shared peak
